# Titin-dependent biomechanical feedback tailors sarcomeres to specialised muscle functions in insects

**DOI:** 10.1101/2024.09.30.615857

**Authors:** Vincent Loreau, Wouter Koolhaas, Eunice HoYee Chan, Paul De Boissier, Nicolas Brouilly, Sabina Avosani, Aditya Sane, Christophe Pitaval, Stefanie Reiter, Nuno Miguel Luis, Pierre Mangeol, Anne C. von Philipsborn, Jean-François Rupprecht, Dirk Görlich, Bianca H. Habermann, Frank Schnorrer

**Affiliations:** Aix Marseille University, CNRS, IBDM, Turing Centre for Living Systems, Marseille, France; Max Planck Institute of Biochemistry, Martinsried, Germany; Department of Neuroscience and Movement Science, Medicine Section, University of Fribourg, Fribourg, Switzerland; Max Planck Institute for Multidisciplinary Sciences, Göttingen, Germany; Aix Marseille University, CNRS, CPT, Turing Centre for Living Systems, Marseille, France

## Abstract

Sarcomeres are the contractile units of muscles that enable animals to move. Insect muscles are remarkable examples because they use extremely different contraction frequencies (ranging from ∼1 to 1000 Hz) and amplitudes for flying, walking and crawling. This is puzzling because sarcomeres are built from essentially the same actin-myosin components. We show here that the giant protein titin is the key to this functional specialisation. I-band titin spans and determines the length of the sarcomeric I-band, and occurs in muscle-type-specific isoforms. Surprisingly, it also rules the length of the force-generating myosin filament in a force feedback mechanism, even though it is not present there. We provide evidence for this model and its validity beyond insects.

**Summary:** Here we identified a mechanical mechanism that instructs sarcomeres to fulfill the specific needs of different muscle types.

## Main Text

Sarcomeres are the basic contractile units of striated muscles across the animal kingdom. They display a stereotypic architecture: regularly arrayed actin filaments are anchored with their plus ends at the Z-discs, central bipolar myosin filaments are linked at a M-band, and both filaments are connected by large titin molecules (*1–4*). The myosin part of the sarcomere, called the A-band, maintains its length during sarcomere contraction, while the myosin-free zone, called the I-band, shortens during contraction (*5*). Despite this conserved architecture, muscle function, as determined by its specific contraction frequency, dimension and force production, varies dramatically across animal evolution. Mammalian muscles generally contract with low Hertz frequencies, shorten about 30% of their length and produce an average power of 5-15 W/kg for the heart or 20-30 W/kg for skeletal muscles during continuous blood pumping or endurance movements, respectively (*5–7*). In contrast, invertebrate muscles achieve a much wider range of contraction regimes: insect flight muscles contract with hundreds of Hertz (up to 1000 Hz in small midges (*8*)), while only shortening 1-2% to produce a mechanical power of more than 80 W/kg to achieve long-range flight (*9–11*). In the same insect, leg muscles sarcomeres contract with low Hertz and shorten up to 50% to power walking or mating. At the extreme end of the evolutionary specialisation are the mandible muscles of leafcutter ants, which produce the highest forces per body mass (>25,000 N/kg body weight to cut leaves) with specialised long sarcomeres (*12*, *13*) and the gut muscles from annelids (Lophotrochozoans) with a sarcomere length of up to 40 µm (*14*). How can evolution tune the muscle parameters to such extremes in a reproducible manner, while maintaining the basic periodic architecture of the contractile sarcomeres?

In mammalian muscles, titin acts as the blueprint for sarcomere architecture: with its N-terminus bound to α-actinin at the Z-disc and its C-terminus embedded at the M-band. Thus, two titin molecules span each sarcomere, which led to the titin ruler hypothesis: the length of the sarcomere and of its myosin filaments are directly controlled by titin protein length (*2*, *3*, *15–17*). In the I-band region, titin contains long PEVK-rich sequences, which act as molecular springs and change length during the sarcomere contraction-relaxation cycles, depending on the pulling forces on the titin protein. These forces are called “passive forces” and are produced by the antagonist muscle that stretches the relaxing muscle (*3*, *18*, *19*). Alternative splicing of titin’s I-band spring region, which removes its PEVK-rich sequence only in the heart, results in 2 µm long sarcomeres with shorter I-bands in the heart and 3 µm long sarcomeres with longer I-bands in human skeletal muscles (*3*, *20–22*). In both types, the myosin filament length is fixed to 1.6 µm by the constant A-band part of titin (*16*). These fixed dimensions of mammalian sarcomeres likely limit muscle specialisations. In invertebrates, sarcomere functions vary much more dramatically, however the mechanisms underlying these morphological and mechanical sarcomere specialisations are not yet known.

### A titin evolutionary tree

To better understand how insects can build sarcomeres with vastly different functional properties from similar components, we used *Drosophila* as a genetically tractable model. We focused on two representative muscle types; first, the larval muscles, which house 8 µm long relaxed sarcomeres with only a small (less than 1 µm) actomyosin overlap, and second the indirect flight muscles, which contain 3.4 µm long sarcomeres with more than 95% actomyosin overlap (*23*, *24*). These specialisations coincide with an alternative splicing of the I-band titin homolog Sallimus (Sls): short Sls correlates with short (200 nm) I-bands in flight muscles, long Sls correlates with long (5 µm) I-bands in relaxed larval muscles (*23*, *25*, *26*). The second titin homolog, Projectin is located selectively at the A-band, which also varies in length between flight and larval muscles (*23*, *25*). Thus, we first wanted to investigated if the presence of two titin homologs in invertebrates instead of one in mammals may explain the variety of muscle specialisations in invertebrates.

To do so, we generated an evolutionary tree of all titin-like genes from humans to jellyfish, extending on previous studies (*27*). As expected, we found one long titin homolog in all chordates, which contains well-known features, including a series of Ig domains and flexible PEVK-rich parts in its I-band part, as well as Ig-Fn3 domain super-repeats in its A-band part, with a kinase domain towards the C-terminal end (Fig. 1A). Interestingly, all Ecdysozoa, including insects and nematodes, contain two distinct titin homologs. One I-band titin, called Sallimus (Sls) in *Drosophila* and ‘Titin homolog’ in *C. elegans*, with the typical Ig and PEVK-rich regions, and one A-band titin, called Projectin (*bt*) in *Drosophila* and Twitchin in *C. elegans*, consisting of the Ig-Fn3 super-repeats and a C-terminal kinase domain (Fig. 1A, B). Lophotrochozoans, including molluscs and annelids, also contain two titins, however in these, the I-band and A-band features appear more mixed (Fig. 1A). One potential titin precursor is also found in jellyfish genomes (Fig. 1A). Thus, based on currently available data, the genomes in the largest part of the animal tree contain two titin homologs, which seem to share the functions of the one mammalian titin.

**Figure 1.**
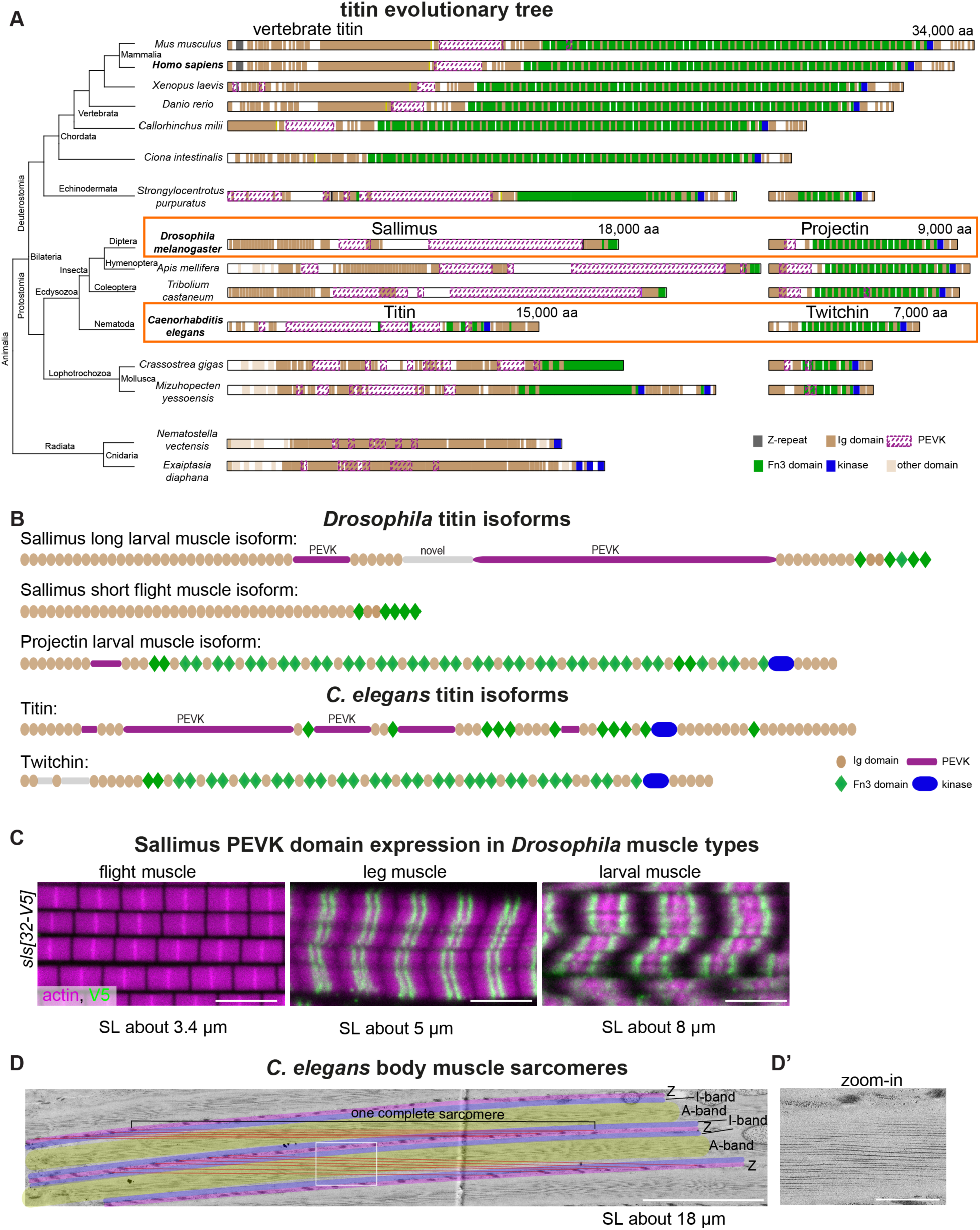
Evolution of titin homologs in animals. (A) Left: Evolutionary tree representing animals with striated muscles. Right: Titin homologs (longest isoforms) are drawn to scale, with highlighted Z-repeats (grey), Ig domains (brown), PEVK (dashed magenta), Fn3 domains (green), and kinase domain (blue). All chordates possess a long titin protein with I-band and A-band features, whereas Ecdysozoa species have two titin homologs separating I-band and A-band features into two proteins. The sizes of the mouse titin and the two *Drosophila* titins (Sallimus and Projectin) and the two *C. elegans* titins (Titin and Twitchin) are indicated. The latter are highlighted by orange boxes. (B) Domain scheme of *Drosophila* and *C. elegans* titin isoforms: Sallimus (short and long), Projectin, *C. elegans* Titin and Twitchin. PEVK-rich regions are highlighted in magenta. (C) Flight, leg and larval muscles of *sls[32-V5]* stained for actin (magenta) and V5 (green). Scale bars: 5 µm. (D) Electron microscopy image of a sectioned adult *C. elegans* body muscle cut in the plane of a sarcomere. For better orientation, Z-disc equivalents, called dense bodies, are overlayed in magenta, I-bands in purple and A-bands in yellow. Entire sarcomeres from one dense body to the next were labelled with red lines. (D’) Zoom-in of indicated white square in D. Scale bars are 5 µm in D and 1 µm in D’.

### Long fly sarcomeres express long titins

To investigate the function of *Drosophila* titins, we focused on the I-band titin Sls, with the rationale that *sls* mRNAs are alternatively spliced, displaying large length differences, while the alternative splicing of Projectin is only minor (*24*). The exons coding for spring-like PEVK-rich sequences are spliced out in flight muscle *sls* isoforms, while they are partially retained in leg muscles and fully retained in larval muscle *sls* isoforms (fig. S1A). Hence, the predicted proteins do contain large PEVK parts in the larval isoform but not in the flight muscle isoform (Fig. 1B). To confirm these mRNA data, we inserted V5 tags into the exons 31 or 32 of the endogenous *sls* gene (see Methods), which code for large parts of the Sls PEVK-rich sequence, generating *sls[31-V5]* and *sls[32-V5]* (fig. S1B and fig. S2). Staining the different muscle types indeed confirmed that the short indirect flight muscle sarcomeres (about 3.4 µm) do not express these *sls* PEVK exons, whereas leg muscle sarcomeres (about 5 µm) and long larval muscle sarcomeres (about 8 µm) do (Fig. 1C and fig. S3a). This demonstrates that sarcomere length and also I-band length (see (*23*, *25*)) indeed correlate with the presence of the flexible spring domains of *Drosophila* I-band titin.

### I-band Titin length impacts sarcomere length

We next asked if the presence of the spring domains in Sls is instructive for sarcomere length and I-band length, as was shown for mammalian titin (*21*). To answer this question, we utilised the fact that our *sls[31-V5]* and *sls[32-V5]* not only inserted a V5-tag but replaced the entire PEVK exon 31 or 32 with the V5-tag. Additionally, we deleted exon 31 or 32 entirely, by replacing them individually with an FRT site, resulting in *sls[31-FRT]* and *sls[32-FRT]*. Finally, we used *sls[31-FRT]* and *sls[32-FRT]* to generate *sls[Δ31-32]*, a *sls* allele with both PEVK exons deleted (fig. S1B and fig. S2). The CRISPR-generated *sls[31-dsRed]* and *sls[32-dsRed]*, which contain a transcriptional STOP, likely resulting in truncated Sls proteins, show the expected early larval lethality with severe sarcomere phenotypes (fig. S3B, C). All other PEVK *sls* deletion alleles were homozygous viable and able to fly, showing that these Sls proteins are functional (fig. S3B).

To investigate sarcomere morphology in the shortened PEVK spring *sls* alleles, we stained relaxed larval muscles with phalloidin to visualise the actin filaments (Fig. 2A). Since we found a correlation between larval size and sarcomere length, we only analysed L3 larval ventral-longitudinal muscles with a length between 400 and 500 µm (fig. S3D, see Methods). Quantifying relaxed sarcomere length showed a length of 8.5 µm in wild-type larval muscles, which is not changed in *sls[31-V5]*. However, sarcomere length in *sls[32-V5]* and *sls[Δ31-32]* muscles is reduced by about 1.5 µm (Fig. 2A, B). The reduced length is compensated by a larger number of sarcomeres in these mutants. This demonstrates that reducing the PEVK spring in Sls does indeed result in shorter sarcomeres.

**Figure 2.**
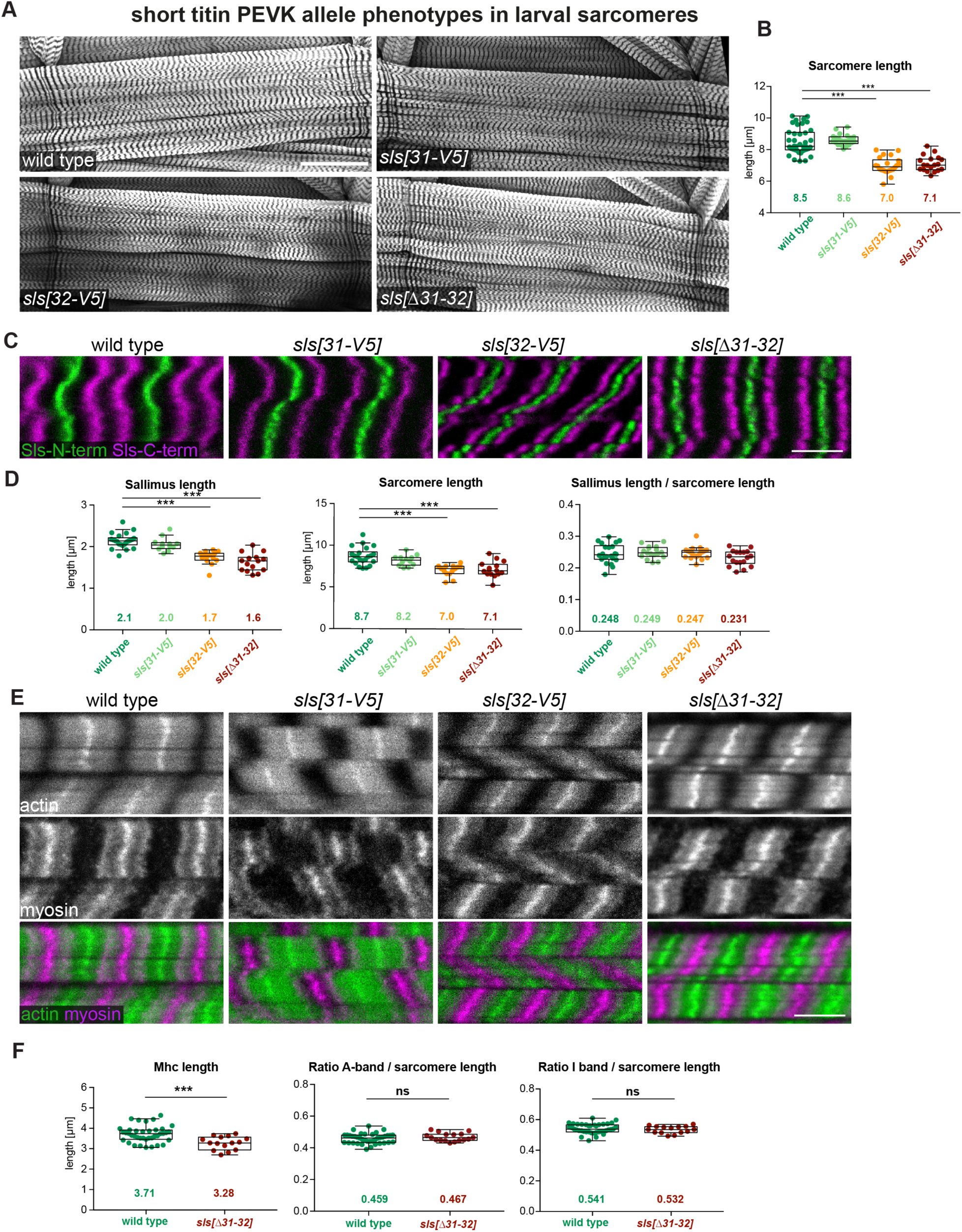
Reducing long titin PEVK length causes shorter larval sarcomeres. (A) Wild-type (*w[1118]*); *sls[31-V5]; sls[32-V5]* and *sls[Δ31-32]* third instar larval VL3 muscles stained for actin (phalloidin). Scale bar: 100 µm. (B) Quantification of sarcomere length from muscle images as shown in (A). Average values are noted in the graphs. Note the shorter sarcomeres in *sls[32-V5]* and *sls[Δ31-32]*. Tukey’s multiple comparisons test, ***: p<0.001. (C) Wild-type (*w[1118]*); *sls[31-V5]; sls[32-V5]* and *sls[Δ31-32]* third instar larval VL3 muscles stained with N- and C-terminal anti-Sls nanobodies (Sls-Nano2 in green and Sls-Nano42 in magenta). Scale bar: 5 µm. (D) Quantification of Sallimus length, sarcomere length and ratio Sls / sarcomere length from muscle images as shown in (C). Tukey’s multiple comparisons test, ***: p<0.001. (E) Wild-type (*w[1118]*); *sls[31-V5]; sls[32-V5]* and *sls[Δ31-32]* third instar larval VL3 muscles stained for actin (phalloidin, green) and myosin (anti-Mhc, magenta). Scale bar: 5 µm. (F) Quantification of myosin filament length (A-band length), ratio A-band / sarcomere length and I-band / sarcomere length from muscle images as shown in (E). Mann Whitney test, ***: p< 0.001, ns: p>0.05.

To directly measure Sls length, we stained larval muscles with a nanobody recognising a domain close to the Sls N-term and one close to its C-term (*23*). We found that Sls in relaxed wild-type larval muscles is about 2.1 µm long, which does not change significantly in *sls[31-V5]*, demonstrating that the PEVK part encoded in exon 31 does not significantly contribute to Sls length in relaxed larval sarcomeres (Fig. 2D). However, Sls length in *sls[32-V5]* and *sls[Δ31-32]* muscles is reduced to about 1.6 µm (Fig. 2D). This reduction of 0.5 µm fits approximately the length of the PEVK domain coded in exon 32 (fig. S3E). These results are very surprising: following the titin ruler hypothesis (*21*), these PEVK deletions should reduce the sarcomere length by subtracting twice 0.5 µm, so 1 µm, compared to wild-type. However, the sarcomere length is reduced by 1.5 µm in *sls[32-V5]* and *sls[Δ31-32]*. Interestingly, we found that the ratio between Sls protein length (which is the I-band length) and sarcomere length scales the same in wild-type and *sls* PEVK mutant sarcomeres (Figure 2D). Together, these data show that Sls length does indeed control the I-band length of *Drosophila* larval sarcomeres, however, the regulation of the entire sarcomere appears to be more complex.

### Titin elasticity controls myosin filament length

To determine the length of the second key filament of sarcomeres, the myosin filament, we stained larval muscles of wild type and *sls* PEVK deletions with a myosin antibody (Fig. 2E) and quantified myosin filament length with an automated tool, PatternJ (*28*), see Methods. We found that myosin filament length was reduced by about 0.5 µm in *sls[Δ31-32]* compared to wild type (Fig. 2F). We confirmed this result by crossing the live myosin filament marker Myofilin-GFP into the *sls[Δ31-32]* background and found a similar difference compared to wild type (fig. S3F). Thus, not only the I-band length scales with sarcomere length but also the A-band length, as the ratio between the A-band and sarcomere length stays constant in wild type and *sls[Δ31-32]* (Fig. 2F). These data were unexpected, as they show that titin’s PEVK-dependent elasticity in the I-band also controls the A-band and hence myosin filament length. This is an interesting finding because it demonstrates that titin impacts sarcomere size in insects, hinting at a leverage mechanism that allows for much larger variations than the protein length itself.

### An artificially elongated titin spring elongates the sarcomere in a muscle-type-specific way

To further explore if changing the length of I-band titin feeds back on the length of the A-band in *Drosophila* muscles, we constructed a fly strain expressing an artificially long Sls protein. To achieve this, we inserted about 4 kb of the PEVK-rich *sls* exon 32, whose deletion caused the above-shown reduction in I-band length, into exon 23 that is present in all *sls* isoforms, resulting in *sls[23-extraPEVK-YPet]* (fig. S4, see Methods). As a control, we only inserted a YPet marker. Adding the extra PEVK spring to Sls results in a further elongation of the already long relaxed larval sarcomeres by about 1 µm (Fig. 3A, B), which is in line with the above-reported reduction in larval sarcomere length in *sls[Δ31-32]*. This is accompanied by the expected gain of Sls length in *sls[23-extraPEVK-YPet]* of about 0.4 µm. Interestingly, in the longer *sls[23-extraPEVK-YPet]* sarcomeres, the myosin filament length does increase as well (Fig. 3C, D). Hence, even long sarcomeres can be further elongated when lengthening the I-band ruler and in turn the A-band scales with it.

**Figure 3.**
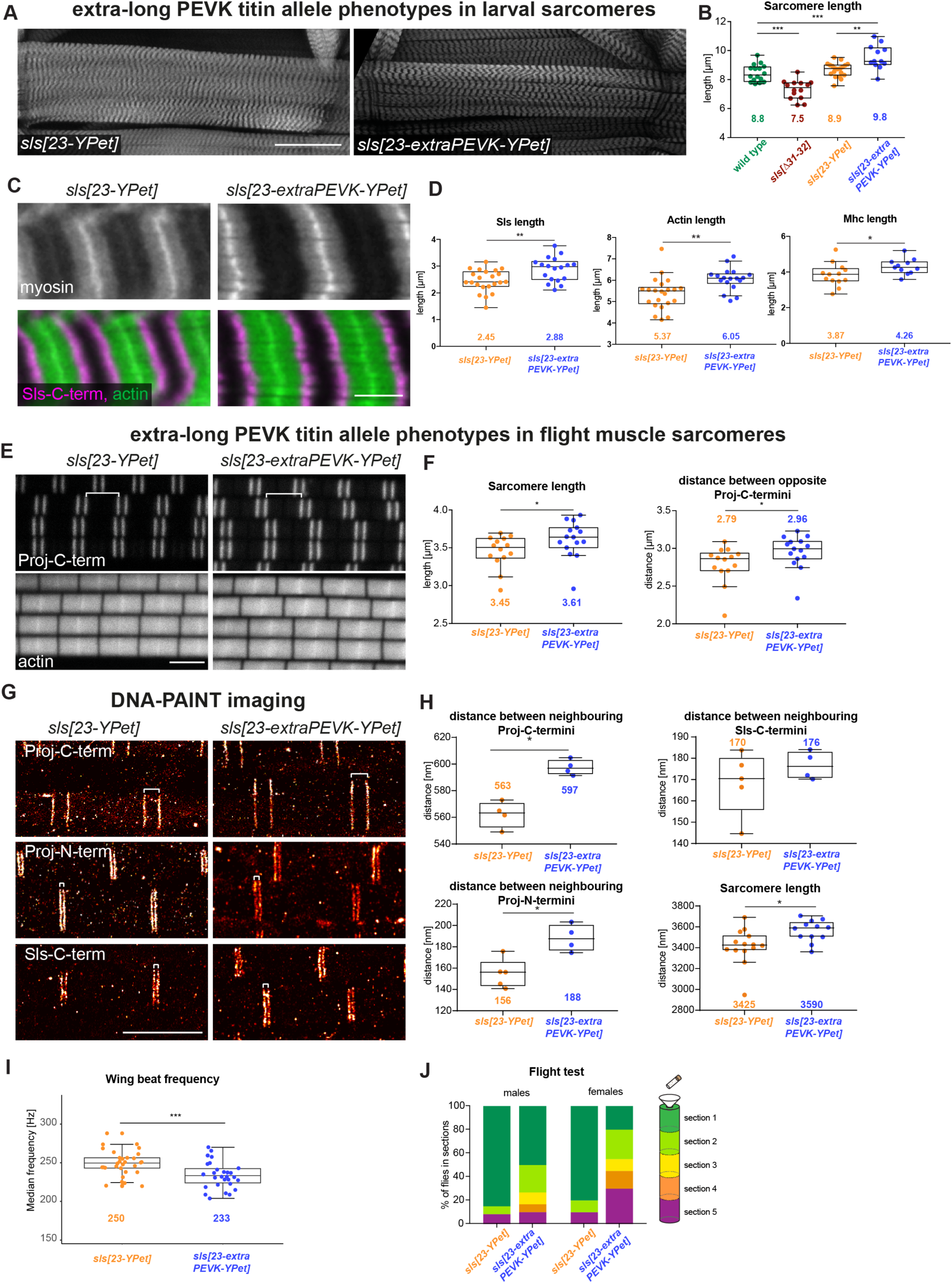
Extra titin PEVK elongates thin and thick filaments in a muscle type-specific way. (A) *sls[23-YPet]* and *sls[23-extraPEVK-YPet]* third instar larval VL1 muscles stained for actin. Scale bar: 100 µm. (B) Quantification of sarcomere length of wild type (*w[1118]*), *sls[Δ31-32]*, *sls[23-YPet]* and *sls[23-extraPEVK-YPet].* Note the longer sarcomeres in *sls[23-extraPEVK-YPet].* Tukey’s multiple comparisons test, *: p<0.05, **: p<0.01, ***: p<0.001. (C) *sls[23-YPet]* and *sls[23-extraPEVK-YPet]* third instar larval VL1 muscles stained for myosin, actin (green) and Sls-C-term nanobody (Sls-Nano48, magenta). Scale bar: 5 µm. Mann Whitney test, *: p< 0.05, **: p< 0.01 ***: p< 0.001. (D) Quantification of Sallimus length, actin and myosin filament length from muscle images as shown in (C). (E) Flight muscles of *sls[23-YPet]* and *sls[23-extraPEVK-YPet]* stained with C-terminal anti-Projectin nanobody (Proj-Nano37) and actin (phalloidin). Scale bar: 3 µm. (F) Quantification of sarcomere length and distance between opposite Proj-C-termini (shown by a white bracket in (E)). Mann Whitney test, *: p< 0.05. (G) DNA PAINT imaging of *sls[23-YPet]* and *sls[23-extraPEVK-YPet]* stained with either anti-Projectin nanobodies (N- or C-terminus, Proj-Nano29 and Proj-Nano37, respectively) or anti-Sallimus nanobodies (C-terminus, Sls-Nano39). (H) Histogram of distances between bands centred around Z-discs. Note the elongation of about 30 - 40 nm. (I) Wing beat frequency quantification of tethered adult *sls[23-YPet]* and *sls[23-extraPEVK-YPet]* flies. Mann Whitney test, ***: p< 0.001. (J) Adult *sls[23-YPet]* and *sls[23-extraPEVK-YPet]* flies tested for flight capability; note the reduced flight ability in *sls[23-extraPEVK-YPet]*. Chi-square test p<0.001.

Since wild-type flight muscles do not express any Sls PEVK sequences (see Fig. 1C, fig. S3a), we predicted that the flight muscle sarcomeres of *sls[23-extraPEVK-YPet]* should also be longer. However, we were surprised to find only a very small sarcomere length increase from about 3.45 µm in control to 3.61 µm in *sls[23-extraPEVK-YPet]* flight muscles (Fig. 3A, B), in contrast to the 1 µm increase in larval muscles. Even more surprising is that this moderate increase is largely due to an increase in myosin filament length, which we assessed by measuring the distance between two opposite Projectin C-termini (Fig. 3E, F) that label the myosin filament about 250 nm before its ends and hence can be resolved by confocal microscopy (*25*).

To directly measure the I-band length with super-resolution microscopy, we quantified Sls and Projectin ends by combining oligo-nucleotide-labelled nanobodies with DNA-PAINT, a technique that achieves about 5 nm spatial resolution in flight muscle tissue (*25*). This not only verified the increased sarcomere length in *sls[23-extraPEVK-YPet]* but also showed that Projectin N- and C-terms are both only about 20 nm further away from the Z-discs compared to controls (Fig. 3G, H), and hence the total I-band length increases only by twice 20 nm. This is also supported by directly measuring the position of the C-terminal end of Sls (Fig. 3G, H). Together, these data show that adding the same extra spring to the I-band titin Sls has muscle-type specific effects: it elongates the larval muscle sarcomeres by 1 µm but the flight muscles sarcomeres by only 120 nm. In both cases, the A-band rescales to the changes in I-band length.

Does this change in sarcomere length have functional consequences? *Drosophila* flight muscles oscillate faster than 200 Hz during flight, while sarcomeres shorten less than 2% (*29*). This requires a very stiff muscle to respond mechanically very quickly. Adding a PEVK spring to Sls in flight muscles should make the sarcomeres more elastic; thus the longer I-band should lower the resonance frequency. To quantify wing beat frequency, we recorded the sound generated from *sls[23-YPet]* and *sls[23-extraPEVK-YPet]* flies during tethered flight (fig. S5A, see Methods). This revealed a lower wing beat frequency in *sls[23-extraPEVK-YPet]* males compared to *sls[23-YPet]* control (Fig. 3I, fig. S5B). To test the consequences of lower wing beat frequency, we performed flight assays and found that flight capability is impaired in *sls[23-extraPEVK-YPet]* males and females (Fig. 3J). This provides a functional explanation for why *Drosophila* flight muscles do contain a short and stiff version of I-band titin Sls.

### Forces across titin correlate with sarcomere length in a muscle type-specific way

Why does inserting the same extra PEVK part of Sls result in sarcomere type-specific effects on Sls length? This hints at important muscle type-specific differences in the forces across Sls. To obtain a first estimate of molecular forces across muscles, we used established force sensors in the integrin adaptor protein Talin in larval muscles, allowing us to measure forces at the muscle-tendon junction (*30*). Comparing control and intra-molecular force sensors, we found that about 30 % of the Talin molecules experience forces above 10 pN in relaxed larval muscles (fig. S6A, B). These forces are not changed in *sls[Δ31-32]*. However, they are larger compared to forces across Talin at adult flight muscle attachment sites (*30*), which suggests that passive stretching forces in relaxed larval sarcomeres are indeed higher compared to flight muscles.

To directly quantify the stretching forces across I-band titin, we inserted these FRET-calibrated molecular force sensors into exon 23 and exon 32 of Sls, generating *sls[23-TS]*, *sls[23-stTS]* and *sls[32-TS]*, *sls[32-stTS]*, respectively, as well as the controls *sls[23-mCherry]*, *sls[23-YPet]*, *sls[32-mCherry]* and *sls[32-YPet]* (fig. S1B, fig. S4C, see Methods). All *sls* sensor flies are homozygous viable, demonstrating the functionality of the internally tagged Sls proteins.

As expected from the mRNA expression data (fig. S1A), all sensors and controls inserted in *sls* exon 23 are expressed in all muscle types, while the ones in exon 32 are not expressed in flight muscles (Fig. 4A). Since we replaced *sls* exon 32 (containing a large PEVK region) with the sensor, the larval sarcomere length of *sls[32-YPet]* is reduced by 1.5 µm, similar to what we had found in *sls[32-V5]* and *sls[Δ31-32]* (Fig. 4B).

**Figure 4.**
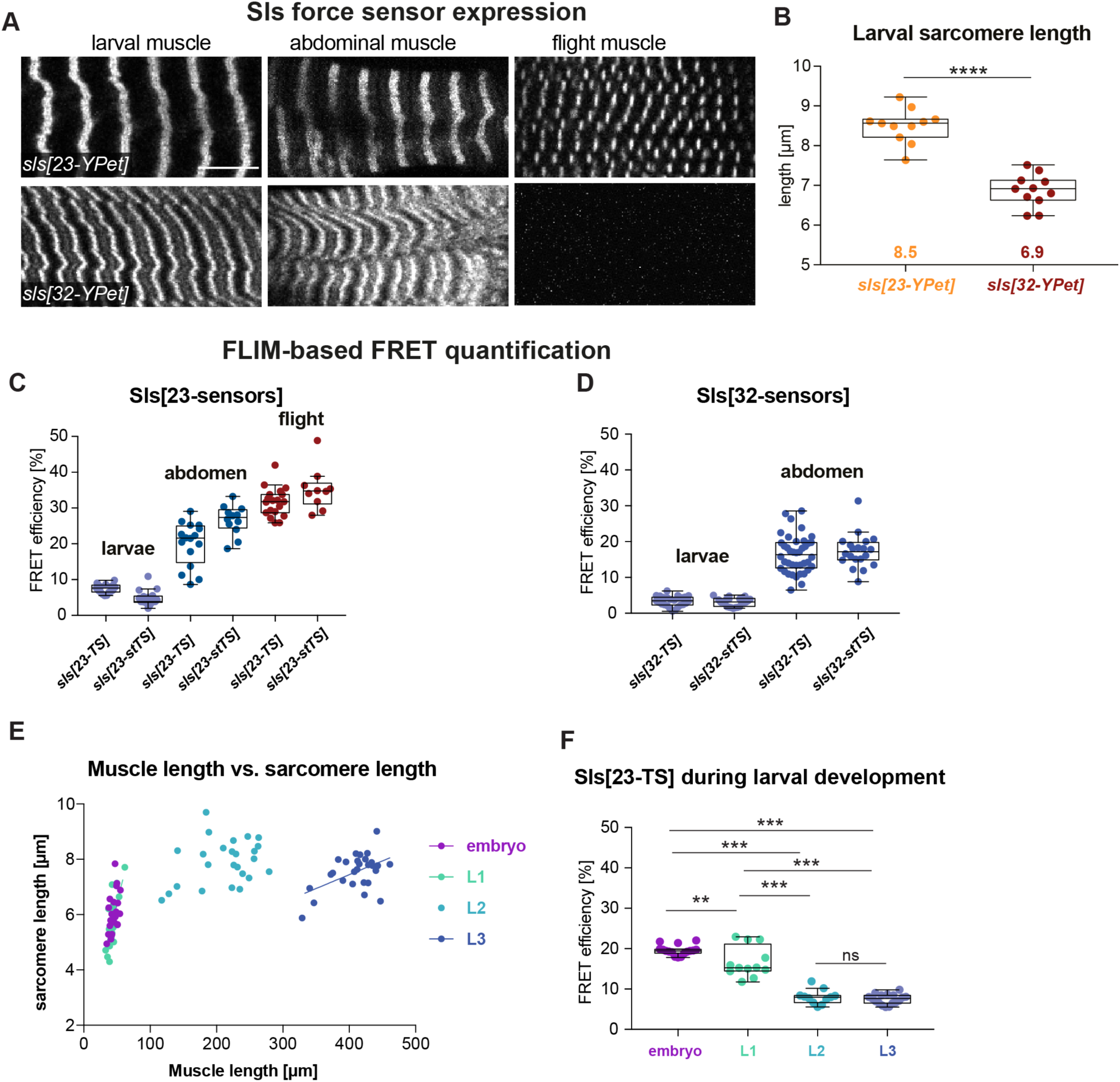
Sallimus molecular forces in different muscle types *in vivo*. (A) *In vivo* expression of *sls[23-YPet]* and *sls[32-YPet]* in third instar larval, adult abdominal and adult flight muscles. Scale bar: 10 µm. Note that *sls[32-YPet]* is not expressed in flight muscles. (B) Quantification of sarcomere length in VL3 muscles of *sls[23-YPet]* and *sls[32-YPet].* Mann Whitney test, ****: p< 0.0001. (C-E) FLIM-based FRET quantification of *sls[23-TS]* and *sls[23-stTS]* (C), and *sls[32-TS]* and *sls[32-stTS]* (D) in larval, abdominal or flight muscles. (E) Relation between sarcomere length and muscle length in Obscurin-GFP (Unc89-GFP) expressing embryos and larvae at different larval instars. (F) FLIM-based FRET quantification of *sls[23-TS]* in embryos and different larval instars. Tukey’s multiple comparisons test, **: p<0.01, ***: p<0.001.

To better characterise these Sls FRET tension sensors, we first measured intermolecular FRET by quantifying the fluorescence lifetime of the YPet donor in the presence or absence of the respective Cherry acceptor in different Sls molecules in trans. We found that intermolecular FRET is negligible (fig. S6C, D). Hence, we measured intramolecular FRET using *sls[23-TS]* and *sls[23-stTS]* in the different muscle types and found very high FRET values in flight muscles, comparable to the no-force control values in the Talin sensor (*30*), suggesting that stretching forces across Sls (the molecular tension) in resting flight muscles are lower than 8 pN and the sensor does not open, thus FRET is high (Fig. 4C). Interestingly, the same sensors display very low FRET in larval muscles and intermediate FRET in abdominal muscles suggesting large stretching forces (tension >12 pN) across Sls in relaxed larval and intermediate forces in the abdominal muscles (Fig. 4C). A similar difference is found in *sls[32-TS]* and *sls[32-stTS]* in larval versus abdominal muscles (Fig. 4D). This is consistent with the hypothesis that long I-band sarcomeres experience high passive forces, stretching Sls to more than 2 µm length. High Sls stretching forces are also supported by the above finding that the deletion of 1,700 amino acids, coded in the PEVK-rich exon 32 of *sls,* reduces Sls protein length by about 0.5 µm, consistent with a fully unfolded conformation of the PEVK-rich sequence (contour length per amino acid: about 0.35 nm (*31*); 1,700 x 0.35 nm = 595 nm).

To investigate if such large forces are already present when the larval muscles assemble sarcomeres at the end of embryogenesis (*23*), we measured FRET across *sls[23-TS]* in stage 17 embryos, as well as first and second instar larvae. We indeed find higher FRET values in embryos and L1 larvae, indicating lower forces, which interestingly correlate with their shorter sarcomere length (Fig. 4F, G). These findings support a novel force-feedback mechanism, in which forces across Sls scale the I-band, which in turn scale the A-band, and thus control the dimensions of the sarcomere.

### A biomechanical feedback model explains sarcomere filament length control

To gain a deeper mechanistic understanding of the molecular mechanism of the feedback, we propose a simple mathematical model that quantitatively explains the scaling of the myosin filament length to that of the extended titin molecules and thus to forces present in the muscle type (Figure 5A). We consider that the sarcomere undergoes cycles of contraction (of duration *T*_c_) and phases of relaxation (of duration *T*_r_). We express the problem in terms of three characteristic lengths, L_A-band_(t), L_I-band_(t) and L_actin_(t), which correspond to the length of the myosin filaments, titin molecules and actin filaments, respectively. During a contraction phase, the myosin motors compress titin up to an equilibrium length, denoted L_I,min_. Within titin’s linear elasticity range, such length is L_I,min_ = T_motor_/K_titin_, with T_motor_ being the maximal compression stress, and K_titin_ the titin elastic modulus. During a relaxation phase, the I-band relaxes to its long equilibrium length, L_I-band_(t)= L_I,max_. Our model proposes that during phases of muscle relaxation new actin and myosin subunits can be added to the filament ends, resulting in a growth of the filaments. In sarcomeres with long I-bands (high force or long titin), the accessibility of myosin motors to the actin filaments is limited, lowering the probability of contraction. This promotes the recruitment of new actin and myosin subunits to grow the filaments; in turn, once the actin filaments have grown up to a length that exceeds sufficiently that of the I-band, an increasing number of myosin motors can bind; once this number N_motors_ exceeds a critical value N_c_, the sarcomeres spontaneously contract (*32*), which blocks new subunit recruitment. This translates into a maximum actin filament length L_actin_ = L_I,max_+ α, with α = N_c_ l, and l the myosin filament length, beyond which the sarcomere contracts. In turn, such maximum actin filament length caps the maximal length of the myosin filaments, set during the contraction phase at L_A-band_(∞) = L_I,max_ + α - L_I,min_. Using this model, we find that the final length of the A-band (L_A-band_(∞) = L_I,max_ + α - L_I,min_) and the actin filaments (L_actin_(∞) = L_I,max_ + α) scale with the relaxed length of the I-band (L_I,max_) (Fig. 5A). Thus, smaller I-band titin molecules or lower titin forces result in a shorter I-band and in turn instruct a shorter A-band (fig. S7A, B). This supports our above *in vivo* findings.

**Figure 5.**
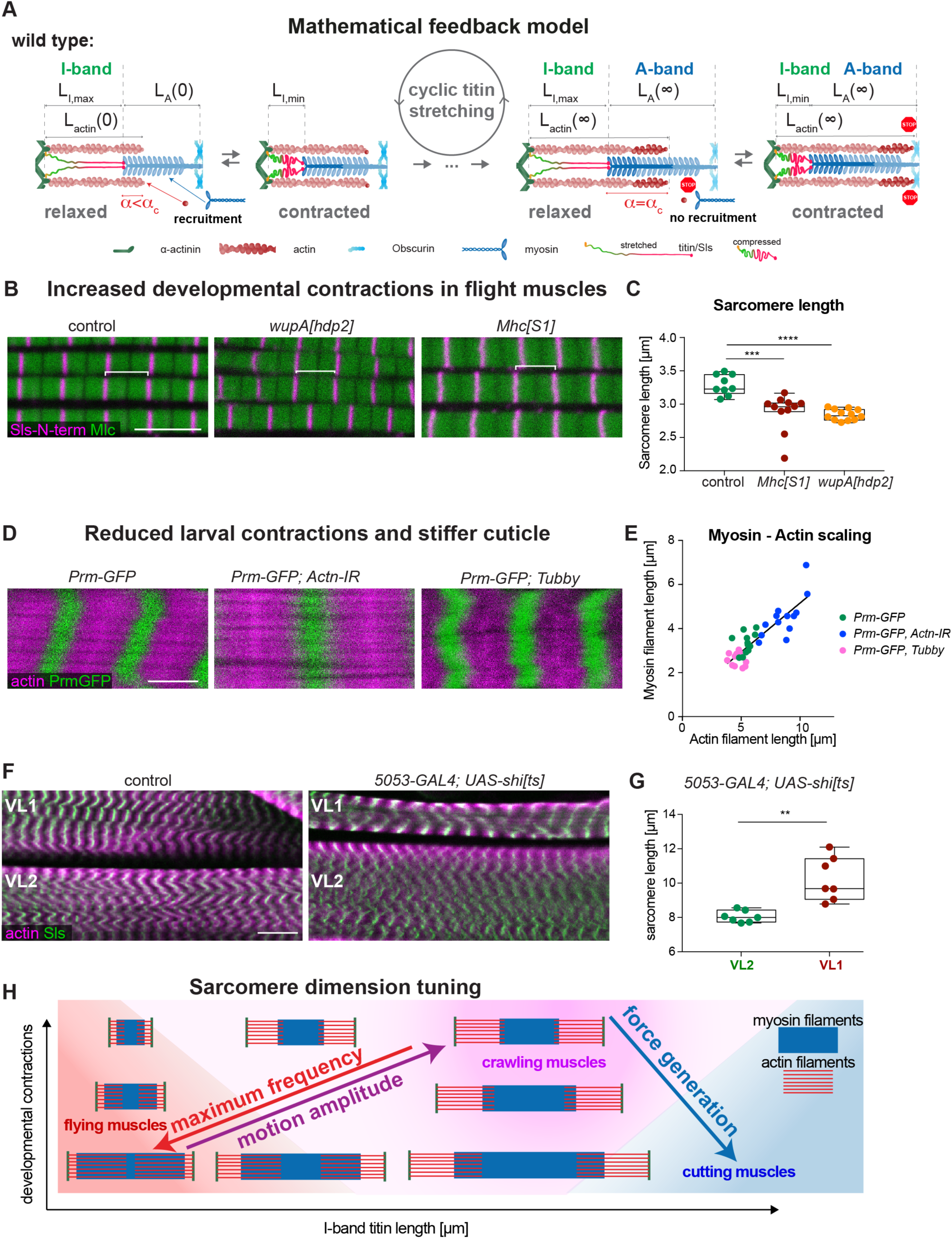
A titin-instructed biomechanical feedback controls filament length. **(A)** Mathematical model of a Sallimus instructed biomechanical feedback regulating I-band and A-band length: actin and myosin filaments grow until a constant actin / myosin overlap (α_C_) is reached in the relaxed state (stretched titin). If in the relaxed state α< α_C_, then actin and myosin filaments can grow by subunit addition at its ends (highlighted by darker colours), as the relaxed state lasts longer. If α= α_C_ is reached in the relaxed state, growth stops as frequent contractions are initiated and recruitment in the contracted state is blocked, because actin bumps into the M-band (red STOP signal). Such, a stable A-band length is reached that scales with the I-band length. See text and Methods for details. **(B)** Wild-type control and hyper-contractile Troponin I (*wupA[hdp2]*) and *Mhc[S1]* mutant flight muscle sarcomeres were stained with Sls-N-term to label the Z-disks (magenta)and an anti-myosin light chain nanobody (NB1-Mlc1) to label the myosin filaments (green). Note the shorter sarcomeres and the shorter myosin filaments in *wupA[hdp2]* and *Mhc[S1]*. Scale bar represents 5 µm. **(C)** Sarcomere length quantification of the data from (B). **(D)** Larval sarcomeres of wild type (*Prm-GFP, Mef2-*GAL4), *Actinin* knockdown (*Prm-GFP, Mef2-* GAL4, *Actn-IR*) or *Prm-GFP Tubby* stained for actin (phalloidin in magenta), and myosin filament length (Prm-GFP, green). Scale bar represents 5 µm. (**E**) Actin and myosin filament length quantification shows the scaling of both. Note that the actin and myosin filaments of the slowly moving *Actinin* knockdown larvae are longer, while in Tubby larvae they are shorter (Mann Whitney test, ***: p< 0.001). (**F**) Larval sarcomeres of wild type and 5053*-GAL4 UAS-shi[ts]* shifted to the restricted temperature at L1 stage. Note the longer sarcomeres of VL1 muscle upon *shi* expression compared to the VL2 control. Scale bar represents 20 µm (**G**). Quantification of F. (**H**) Phase diagram that illustrates how the two variables, developmental contractions and I-band titin length, can generate sarcomeres with vastly different dimensions that are specialised for maximum frequency at high force (flying muscles, red), large motion amplitude (crawling muscles, magenta) or maximum force generation (cutting muscles, blue).

This model predicts that reducing the sarcomere contraction probability (or mathematically increasing α) will increase sarcomere length, as well as the actin and myosin filament length because the time to grow the filaments at the relaxed stage is increased (fig. S7A, C). Interestingly, this scenario is at play during flight muscle development and results in the observed long actin and myosin filaments with 95% overlap despite having a short I-band: at 48 h after puparium formation (APF) sarcomeres are 2 µm short and spontaneously contract. After 48h APF, these contractions stop until 90 h APF when flies eclose with 3.4 µm long flight muscle sarcomeres that have 3.2 µm long myosin filaments (see Fig. 3) (*24*). Interestingly, in various mutants, in which the spontaneous muscle contractions continue after 48 h APF, the sarcomeres remain shorter at 90 h APF (*24*, *33*– *36*). Strikingly, we found that sarcomeres of particular *myosin heavy chain* or Troponin I (*wupA*) mutations, which cause excessive contractions during development (*37*, *38*), are abnormally short, containing shorter myosin filaments (Figure 5B, C). Thus, our force-feedback mechanism can explain how sarcomeres with a maximum actomyosin overlap (a very large α in the model fig. S7A, C), hence powerful sarcomeres that can oscillate at high frequency to mediate insect flight, can be made during development.

To explore if filament growth can also be induced ectopically in sarcomeres that do readily contract, we choose the larval muscles and knocked-down α*-actinin*, which strongly reduces larval locomotion and thus sarcomere contractility (fig. S 7B, C and Video S1). Consistent with our proposed feedback, we found that not only sarcomere length but also actin and myosin filament length is increased in these slowly moving larvae (Fig. 5D, E; fig. S7D-G).

To further challenge the model, we blocked sarcomere contractions in a single muscle cell per larval hemi-segment only, by expressing dominant negative Dynamin (*shibire [ts]*) (*39*) (see Methods). Strikingly, the silenced muscle VL1 displays elongated sarcomeres with longer actin filaments, compared to its direct VL2 neighbour analysed in the same larva (Fig. 5F, G; fig. S7H). This demonstrates that extended periods of muscle relaxation result in an elongation of actin and myosin filaments, as predicted by our model, and hence result in longer sarcomeres without the need for a gigantic titin A-band ruler protein.

Finally, we aimed to mechanically manipulate the muscle independently of the sarcomere components. To do so, we took advantage of a dominant mutation in the *Tubby* gene (*Tb[1]*) that results in short and fat larvae. As Tb is a chitin-modifying enzyme, likely the exoskeleton is more rigid resulting in the short pupae (*40*, *41*). Interestingly, we confirmed that sarcomeres in these Tubby larvae are shorter, as had already been shown (*42*), and we found that both actin and myosin filaments are shorter, too. Strikingly, both filaments scale linearly in all the genotypes in larval muscles (Fig. 5D, E, fig. S7G). This demonstrates that changes in the exoskeleton feedback on the sarcomere components to adjust filament length. This might be an advantage to better adapt to variable endogenous or exogenous conditions during life as well as during evolution to adjust to variable thorax sizes and wing beat frequencies, for example in large butterflies.

Taken together, we have uncovered a biomechanical mechanism to regulate the dimension of the sarcomere in the different *Drosophila* muscle types that is both genetically and mechanically controlled. The length of the I-band titin as well as developmental contractions can tune the filament and sarcomere length to allow the formation of very stiff muscles with an extensive actomyosin overlap, which oscillate at high frequency to power insect flight. In the same organism, much more elastic muscles, which contract slower but at higher amplitude, to support larval crawling and adult walking can be constructed (Fig. 5H). Both would not be compatible with the strict titin ruler that is applied in mammalian sarcomeres.

### Evolutionary conservation

Is the force-feedback mechanism conserved outside of insects? Our titin evolutionary tree revealed two distinct titin homologs not only in beetles but also in nematodes (Figure 1A, B), suggesting that a similar mechanism may control actin and myosin filament length in the sarcomeres of a wide variety of non-vertebrate species.

To investigate the sarcomere structure of the nematode *C. elegans*, we used electron-microscopy. This revealed the unique tilted arrangement of the actin and myosin filaments in *C. elegans* body muscle sarcomeres, which makes length quantifications difficult (*43*). Sectioning adult *C. elegans* in the plane of the myosin filaments revealed a sarcomere length of about 18 µm containing very long (about 15 µm) myosin filaments (Fig. 1D, D’). As the A-band titin homolog Twitchin is only about 7000 amino acids long and, like Projectin in *Drosophila* larval muscles (*23*), decorates the entire myosin filament (*44*), a classical titin ruler mechanism of myosin filament length control is neither possible in nematodes nor in insects. Thus, we have uncovered how the vastly different sarcomere dimensions present in a large clade of the animal tree are set in a controlled way to match the biomechanical requirements of specialised tailor-made muscle types.

### Conclusions

The here revealed force-feedback mechanism controlling the length and architecture of the key sarcomere filaments provides a molecular explanation for how animals can produce tailor-made sarcomeres for specialised muscle functions. By applying this mechanism, sarcomeres tuned for fast oscillations powering insect flight, sarcomeres tailored for large motion amplitudes like walking or larval crawling or highest force production to cut fibrous leaves can be built within the same animal (Figure 5C).

As this covers vastly different sarcomere dimensions, including variations of the myosin filament length, a strict titin ruler mechanism cannot apply, simply because it would need a gigantic A-band titin gene, much larger than the mammalian one. Additionally, this mechanism equips the muscles with more flexibility. It allows an animal to adapt its muscles to changes in their mechanical environment, possibly during ageing, or to changes of the animal’s internal state. One important physiological adaptation occurs in females after mating to effectively produce eggs. Mated females eat more and their gut enlarges in diameter (*45–47*). As a consequence, their circular muscles need to grow in length to cover the larger gut surface. Smartly, *Drosophila* females solve this problem by increasing their actin, myosin and consequently sarcomere length after mating (*Mineo et al, submitted*). This allows females to absorb food more effectively and thus produce more eggs. In summary, the here described force-feedback mechanism allows muscles to specialise and adapt to the particular tasks, which may be one of the key reasons for the success of insects and nematodes during evolution.

## Supporting information

Loreau Supplementary Figures

File S1 - code to calculate the PEVK content

Table S1 - Titin evolutionary tree, species and protein names

## Acknowledgments

We thank Celine Guichard, Katja Finkl and Bettina Stender for fly embryo injections, fly maintenance and excellent technical assistance. We are grateful to Anthony Cammarato, Troy Littleton, the Bloomington and the VDRC stock centers for fly stocks. We would like to thank Stefan Raunser and Mathias Gautel and all their group members as well as the Schnorrer and Görlich group members for their stimulating discussions within the StuDySARCOMERE ERC synergy grant. We are indebted to the IBDM imaging and fly facilities for help with image acquisition, maintenance of the microscopes and production of fly food.

## Author contributions

Conceptualization: FS, VL, WK

Methodology: VL, WK, EHC, PDB, NB, SA, AS, CP, SR, NML, PM, AvP, JFR, DG, BHH, FS

Investigation: VL, WK, EHC, PDB, NB, SA, AS, CP, SR, JFR

Visualization: VL, WK, EHC, PDB, NML, PM, JFR, BHH, FS

Funding acquisition: FS, BHH, DG, JFR, AvP

Supervision: FS, BHH, DG, JFR, AvP

Writing – original draft: FS, VL

Writing – review & editing: FS, DG, VL, EHC, PM, NML

## Data and materials availability

All data are available in the main text or the supplementary materials.

## Supplementary Materials

### Materials and Methods

#### Drosophila strains and genetics

Fly stocks were maintained under standard culture conditions at 27 °C (*48*). All new *sls* alleles generated in this study are listed in Fig. S1B. The *Zasp66-GFP* allele was used to label the Z-disc (*49*), *Prm-GFP* (fTRG-475) to label the A-band (*50*), *Mhc[weepGFP]* to label a subset of Mhc isoforms (*51*), *Talin-YPet*, *Talin-TS* and *Talin-C-TS* to quantify force across Talin (*30*). The CRISPR injections were done into *Act5c-Cas9, DNAlig4[169]* (*30*). Heat-shock flippase was used to generate *sls[Δ31-32]*) and *Df(3L)BSC366* was used as *sls* deficiency (both from Bloomington). The *Actn* RNAi line (V7760) was from VDRC (*52*). The *Mhc[S1]* allele was from Troy Littleton and the *wupA[hdp2]* (Troponin I) allele from Anthony Cammarato (*38*). Canton S or *w[1118]* were wild-type controls.

#### Titin evolutionary tree

Human titin (NP_001243779.1) and *Drosophila melanogaster* Sallimus (NP_001261304.1) and Projectin/bent (NP_001162825.1) were downloaded from the NCBI Protein database and used with BLAST (*53*, *54*) to search for reciprocal best hits (RBH)(*55*) with Deuterostoma and Protostoma, and their subphyla respectively. Because of its length and its repetitive nature, titin may be mis-sequenced or confused with obscurins. Thus, we manually selected only orthologs we found to have strong evidence based on the quality of available sequencing data. Thus, we kept the 15 species presented in Figure 1A. Their protein identifiers are listed in Supplementary Table 1 and the sequences in Supplementary Data 1.

We used InterPro (*56*, *57*) to predict the different domains in each ortholog. Using a Python3 script, we curated the results, keeping only our domains of interest, i.e., Ig, Fn3 and kinase domains and Z-repeats. Even though InterPro can calculate the presence of PPAK motifs, repeated motifs specific to PEVK domains, we calculated the presence of these PEVK domains with a Python3 script which calculates the proportion of P, E, V and K residues in a given window of 250 amino acids. Windows with more than 40% of P, E, V and K were considered as PEVK domains. The Python script is also attached in Supplementary File 1. In the end, the proteins were represented using R with respect to their sizes, domains and PEVK regions. As the orthologs are very different, it was impossible to build a phylogenetic tree. We thus decided to use the common species tree proposed by the NCBI taxonomy (*58*) for visualisation purposes.

#### CRISPR and RMCE

Modification of Sallimus length and the insertion of tension sensors were made by combining CRISPR/Cas9-mediated genome engineering with phiC31-mediated cassette exchange as described (*59*). Briefly, the genomic sequence in the *Act5c-Cas9, DNAlig4[169]* was sequenced, the sgRNAs were designed with an online tool (https://crisprscan.org) and transcribed *in vitro* (MEGAshortscript^TM^ T7 Kit, Invitrogen). The efficiency of the sgRNAs was tested by embryo injection into *Act5c-Cas9* or in Cas9-expressing S2-cells with a T7 endonuclease assay (*59*). For step 1, one pair of sgRNAs (100 ng/µl) was injected into *Act5C-Cas9, DNAlig4[169]* embryos in combination with a dsRed donor vector (500 ng/µl) containing a dsRed eye marker cassette flanked by attP sites and homology arms (see fig. S2 and S4). Successful homologous recombination events were identified by red fluorescent eye colour and verified by sequencing. For step 2, vasa-phiC31 plasmid (200 ng/µL) was injected into step 1 embryos together with an attB-donor vector (150 ng/µL). Successful exchange events were identified by the absence of red fluorescent eyes, and the correct orientation of the cassette was verified by PCR. Homozygous stocks were established from single flies.

#### Flight and leg muscle staining

Detailed flight and leg muscle staining protocols were recently published (*23*, *60*). Briefly, the head, abdomen and wings were clipped from young adult flies (3 - 7 days old) and thoraces were fixed in 4% PFA in PBS-T (PBS with 0.3% Triton X-100) for 20 min at room temperature. After washing once with PBST, thoraxes were placed on a slide with double-sticky tape with the anterior facing the tape and cut sagittally with a microtome blade (Pfm Medical Feather C35). Hemi-thoraces were stained with fluorescent nanobodies and rhodamine-phalloidin (1:1000 Molecular Probes) for 2 hrs at room temperature (RT) or overnight at 4°C. Hemi-thoraces were washed twice with PBS-T, mounted in SlowFadeTM Gold Antifade (Thermofisher) using two coverslips as spacers and flight or leg muscles were imaged with a Zeiss LSM880 confocal microscope using a 63x objective.

#### Dissection and staining of larval muscles

A detailed protocol for staining of larval muscles was published (*23*). Briefly, third instar (L3) larvae were either immobilised by dipping for 1 second in 65°C water (*61*) or alive covered with HL3 buffer (70mM NaCl, 5mM KCl, 1.5mM CaCl_2_, 20mM MgCl_2_, 10mM NaHCO_3_, 5mM Trehalose, 115mM Sucrose, 5mM HEPES)(*23*), pinned individually and dissected with sharp scissors from the dorsal side. Interior organs were removed with forceps and the larval filets were fixed in 4% PFA in PBS-T (PBS with 0.3% Triton X-100) for 30 min and then blocked in 4% normal goat serum for 30 min at room temperature. Nanobodies and antibodies were incubated in PBS-T overnight at 4°C. Larval filets were then washed 3 times 10 min in PBST at RT and stained with secondary antibodies and phalloidin (labelled with rhodamine 1:1000, Molecular Probes) in PBS-T for 2 hrs at RT in the dark. After washing 3 times with PBST for 5 min, larval filets were mounted in SlowFadeTM Gold Antifade (Thermofisher) and imaged with a Zeiss LSM880 confocal microscope using 20x, 40x or 63x objectives.

#### Antibodies and nanobodies

Primary antibodies (anti-Mhc (3e8-3D3, mouse, DSHB, 1:100), anti-V5 (mouse, Invitrogen, 1:1000)) or fluorescently labelled nanobodies (Sls-Nano2 (1:1000), Sls-Nano39 (1:1000), Sls-Nano42 (1:1000), Sls-Nano48 (1:1000), Proj-Nano37 (1:1000) (*23*)) were incubated overnight at 4°C. After washing 3 times 10 min in PBST at RT, samples were stained with secondary antibody (Alexa488 goat anti-mouse IgG, 1:500, Molecular Probes) and/or phalloidin (Rhodamine conjugate, 1/500, Molecular Probes) in PBST for 2 hrs at RT in the dark. The newly made anti-myosin light chain nanobody NB1-Mlc1 was generated by immunising a trimeric complex of Mlc1, Mlc2, and a cognate myosin heavy chain fragment (comprising residues 709-840). This complex was generated by co-expressing the three proteins as His14-ScSUMO and His14-NEDD8 fusions in *E. coli* and purifying them by capture to Ni-chelate beads followed by a tag-cleaving elution with Ulp1 and NEDP1 (*62*). Immune library generation, phage display, nanobody production and nanobody labelling were performed as described (*23*, *63*). Biolayer interferometry measurements revealed that the nanobody targets the Mlc1 subunit with a K_D_ of ∼1nM.

#### Image quantification and filament length measurements

Quantifications of protein or filament lengths and distance between proteins were achieved using the automated algorithms of the Fiji macro toolset PatternJ (*28*, *64*). In brief, after the manual selection of a myofibril in Fiji, the resulting myofibril intensity profile is used by the PatternJ algorithms to extract the position of bands or edges of filaments in the following way:

1. The user specifies to PatternJ the main features of protein stainings in a sarcomere from predefined patterns (one or multiple bands, blocks, blocks with bands).
2. PatternJ obtains the average sarcomere length using the autocorrelation of its intensity profile. Based on the position of the highest intensity in the profile it defines a reference sarcomere.
3. It extracts the position of all sarcomeres using a cross-correlation between the intensity profile of the myofibril and the intensity profile of the reference sarcomere, allowing for the segmentation of each sarcomere.
4. Once each sarcomere is segmented, particular features defined in the first step are then located and localised precisely by fitting with relevant mathematical functions, such as a Gaussian function for a thin band or a spline function to locate the edge of a filament.

Based on the position of the features extracted with PatternJ, we straightforwardly obtained the different lengths of filaments or distances between bands.

#### DNA-PAINT

For DNA-PAINT, adult flies were fixed as described above for immuno-stainings and hemi-thoraces were stained and imaged as described recently (*25*). Briefly, Sls-Nano39, Proj-Nano29 or Proj-Nano37 coupled to P3 oligos (*25*) (about 50 nM) were incubated with fly hemi-thoraces overnight. After washing with PBS, 1% Triton, stained hemi-thoraces were mounted in the imaging chamber and the respective imager oligos coupled with Cy3B (Metabion) were added in imaging buffer, Buffer C (1× PBS, 500 mM NaCl), at 2 nM. Directly before imaging, Buffer C was supplemented with 1× Trolox, 1× PCA and 1× PCD. 100× Trolox: 100 mg Trolox, 430 µl 100% methanol, 345 µl 1 M NaOH in 3.2 ml H_2_O. 40× PCA: 154 mg PCA, 10 ml water and NaOH were mixed, and pH was adjusted to 9.0. 100× PCD: 9.3 mg PCD, 13.3 ml of buffer (100 mM Tris-HCl pH 8, 50 mM KCl, 1 mM EDTA, 50% glycerol). All three were frozen and stored at - 20°C. TIRF imaging of the single molecule DNA hybridisation events was done on a TIRF illumination system (SAFE 360, Abbelight, France) combined with an inverted microscope, using a 100X N.A. 1.47 objective (Evident, Japan). Coverslips with embedded gold particles were used for drift correction (550-100Auf, Hestzig LLC). Images were processed and bands were fitted as described in detail recently(*25*).

#### Behavioural tests

Tethering and flight frequency recording: male flies were anaesthetized on ice, then moved to a Peltier-cooled aluminium block kept at 4° C and fixed with their notum onto a tungsten pin with UV hardening glue (Tetric EvoFlow, Ivoclar Vivadent), as described previously (*65*, *66*). The wire was then fixed on a block of plasticine to suspend the male (fig. S5A). The tethered fly was provided with a polystyrene ball (diameter 2-3 mm) as support and then placed above an electret condenser microphone (CMP-5247TF-K, CUI Inc) (distance 2 cm). After a recovery of approx. 1 min, the ball was removed with a gentle air puff, triggering flight. The flight was recorded for 10 minutes, amplified with a custom-made circuit board, and digitized with a multifunction data acquisition device (NI USB-6259 MASS Term, National Instruments) (*66*, *67*).

Flight wing beat frequency analysis: analysis of the flight wing beat frequency from recorded signals was performed with Raven 1.6 (The Cornell Lab of Ornithology) using a Fast Fourier Transform (FFT) type Hann, with a window length of 10000 samples and 0% overlap. The sound was filtered with a band-pass filter (between 150 and 350 Hz) to exclude harmonics beside the fundamental band. For each flight bout, the first seconds were excluded from the analysis to avoid possible interference due to the air puff used to trigger flight.

Flight behaviour: to test for flight, about 30 male flies (1 - 3 days old, aged at 25°C) were sorted into a fresh food vial and aged for another 24 h. Then, they were thrown through a funnel at the top of a 1 m × 8 cm plexiglass cylinder with 5 marked sections according to an established protocol (*61*). The landing positions of the flies were noted. Flight assays were performed in triplicates with a total of at least 80-100 males tested.

Larval locomotion: to test larval muscle function, we used a recently published protocol (*23*). Briefly, third instar (L3) larvae were collected at the wandering stage, placed on an apple agar plate and allowed to acclimatise for at least 20 min at room temperature. Larvae were then placed simultaneously in the centre of the plate and imaged at a frame rate of 25 Hz. Images were acquired using an infrared Basler acA2040-90 µm NIR camera equipped with a Kowa LM12SC lens and a homemade LED infrared illumination system (WINGER WEPIR3-S1 IR Power LED Star infrared at 850 nm). The Pylon viewer software from Basler was used to control acquisition, and exposure time was adjusted for enhanced contrast. The assays were repeated at least 3 times for each genotype, with assays done on different days. The videos were analysed using FIMTrack(*68*), and data were visualized via Python.

#### FLIM-FRET quantification

The FLIM data were acquired on Zeiss LSM880 confocal equipped with a PicoQuant LSM Upgrade Kit using a PicoQuant Time-Correlated-Single-Photon detector (TimeHarp 260, in version 250 ps (NANO module) base resolution) and a femtosecond pulsed 510 nm laser running at 100 MHz repetition rate. The data were analysed using the SymPhoTime 64 software (PicoQuant). First, an intensity image was created by manually drawing an ROI around the target structure (Z-discs or muscle attachment sites). Second, photon arrival times of all photons inside the ROI were plotted in a histogram, and the tail of the curve was fitted with a mono-exponential decay fit for the YPet donor alone or with a bi-exponential decay fit for all the different tension sensors to calculate an average fluorescence lifetime τ. Third, the FRET efficiency (E) for each tension sensor was calculated according to the following formula, with τ_DA_ being the lifetime of the donor in the presence of the acceptor (the tension sensor pair) and τ_D_ the lifetime of the donor alone (YPet control):

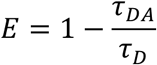

For all measurements, τ_D_ was determined as the median lifetime of either Talin [YPet], Sls[23-YPet] or Sls[32-YPet] using the same experimental conditions. Experiments were repeated more than 3 times on different experiment days.

#### Mathematical modelling

In our simulations, we consider that the sarcomere unit undergoes cycles of contraction (of duration *T*_c_) and phases of relaxation (of duration *T*_r_). The system evolves linearly within each phase, with

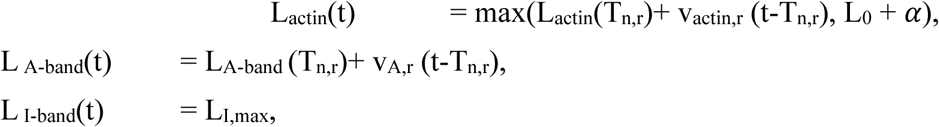

during the n-th relaxation phase, with T_n,r_ the start date of the last relaxation phase, v_r_ the actin and A-band growth speed in the relaxation phase, respectively, and

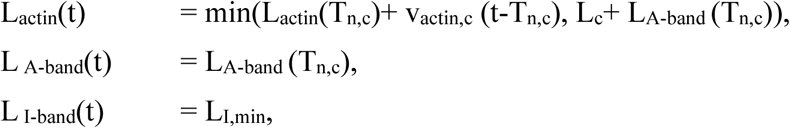

during the n-th contraction phase, with T_n,c_ the date of the start of the last contraction phase. Due to the wall condition at the center of the sarcomere, the actin filaments cannot exceed the sum of the titin and myosin lengths L_actin_(t)< L_I-band_(t) + L_A-band_(t), see Fig. 5A; we make sure that our initial conditions verify this condition. We then solve the system of equations numerically and consider the final state reached. As expected (see main text), both the final lengths of the actin (L_actin_(∞) = L_I,max_ + α) and A-band (L_A-band_(∞) = L_actin_(∞) - L_I,min_ = L_I,max_+ α - L_I,min_) scale with the relaxed length of the I-band (L_I,max_).

Figure S1 – *sallimus* (*sls*) alleles

Figure S2 – *sls[31]* and *sls[32]* CRISPR - RMCE editing

Figure S3 – Sls-V5 expression and *sls* alleles sarcomere phenotypes

Figure S4 – *sls[23]* CRISPR - RMCE editing

Figure S5 – wing beat frequency set-up and recordings

Figure S6 – Talin and Sallimus molecular forces

Figure S7 – Actin-myosin filament length scaling

Table S1 – Titin evolutionary tree, species and protein names.

Video S1 – Tracked wild type and *Mef2-*GAL4, *Actn-IR* larvae.

Data S1 – Archive of all titin protein FASTA sequences used for evolutionary tree in Figure 1.

Data S2 – Data of Figure 2

Data S3 – Data of Figure 3

Data S4 – Data of Figure 4]

Data S5 – Data of Figure 5

File S1 – Python script to calculate the PEVK content in the titin sequences.

