## Supplementary material for "Titin-dependent biomechanical feedback tailors sarcomeres to specialised muscle functions in insects": Loreau Supplementary Figures

**The PDF file includes:**

Figure S1 – *sallimus* (*sls*) alleles

Figure S2 – *sls[31]* and *sls[32]* CRISPR - RMCE editing

Figure S3 – Sls-V5 expression and *sls* alleles sarcomere phenotypes

Figure S4 – *sls[23]* CRISPR - RMCE editing

Figure S5 – wing beat frequency set-up and recordings

Figure S6 – Talin and Sallimus molecular forces

Figure S7 – Actin-myosin filament length scaling

**Other Supplementary Materials for this manuscript include the following:**

Table S1 – Titin evolutionary tree, species and protein names.

Video S1 – Tracked wild type and *Mef2*-GAL4, *Actn-IR* larvae.

Data S1 – Archive of all titin protein FASTA sequences used for evolutionary tree in Figure 1.

Data S2 – Data of Figure 2

Data S3 – Data of Figure 3

Data S4 – Data of Figure 4

Data S5 – Data of Figure 5

File S1 – Python script to calculate the PEVK content in the titin sequences.

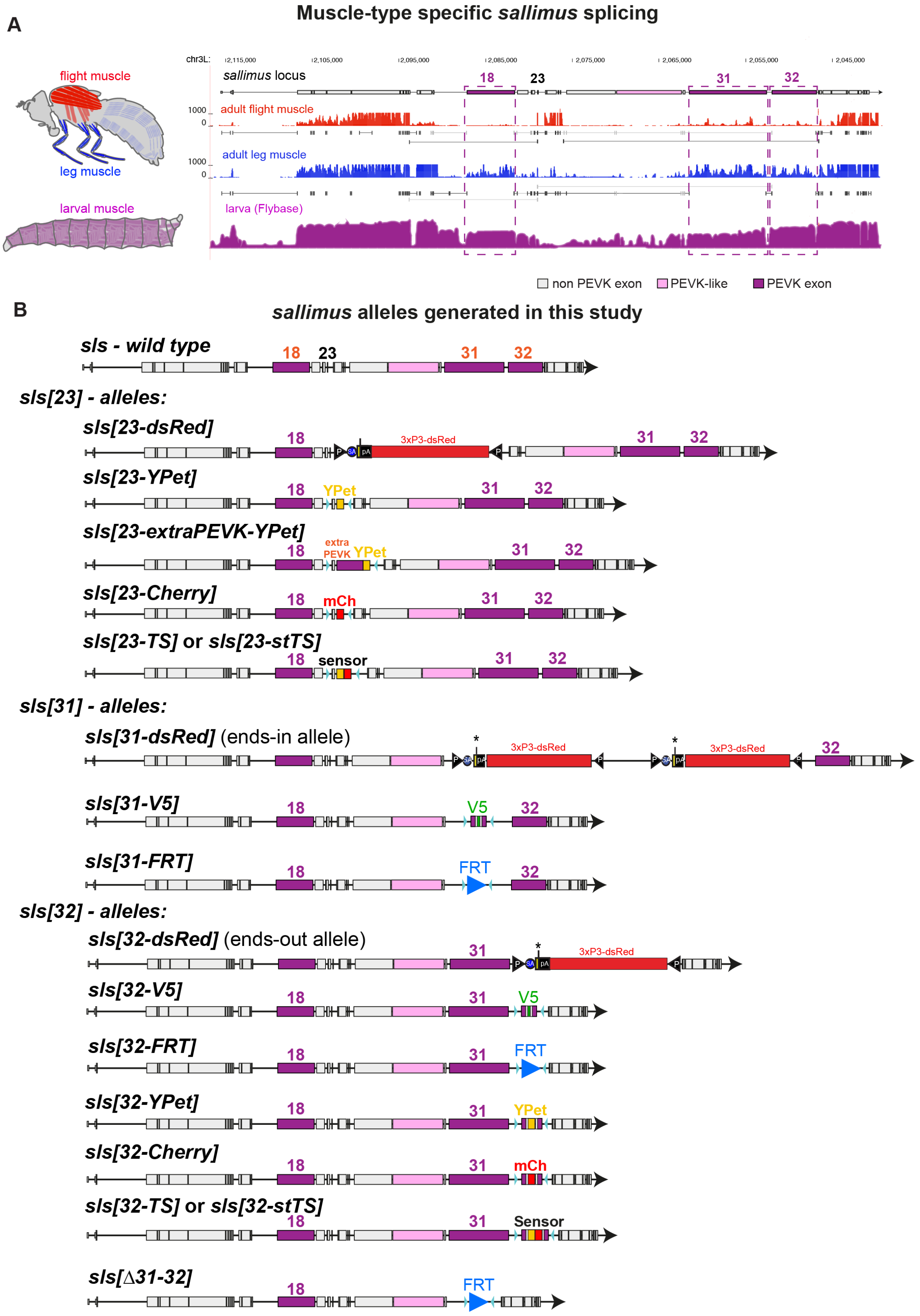
A
Muscle-type specific sallimus splicing
flight muscle
leg muscle
larval muscle
chr3L:
sallimus locus
18
23
31
32
adult flight muscle
adult leg muscle
larva (Flybase)
non PEVK exon
PEVK-like
PEVK exon
B
sallimus alleles generated in this study
sls - wild type
sls[23] - alleles:
sls[23-dsRed]
3xP3-dsRed
sls[23-YPet]
YPet
sls[23-extraPEVK-YPet]
extra PEVK
sls[23-Cherry]
mCh
sls[23-TS] or sls[23-stTS]
sensor
sls[31] - alleles:
sls[31-dsRed] (ends-in allele)
sls[31-V5]
V5
sls[31-FRT]
FRT
sls[32] - alleles:
sls[32-dsRed] (ends-out allele)
sls[32-V5]
sls[32-FRT]
sls[32-YPet]
sls[32-Cherry]
sls[32-TS] or sls[32-stTS]
Sensor
sls[Δ31-32]

**Figure S1**

**Fig. S1 – *sallimus* (*sls*) alleles**

(A) mRNA-SEQ read counts and junction reads (grey lines) in the sallimus locus from dissected adult *Drosophila* indirect flight muscles (red), adult legs (blue) and larvae (magenta). PEVK domains are highlighted in magenta. Data were extracted from (*24*) and Flybase. (B) Scheme of all new *sls* alleles generated in this study; PEVK regions are highlighted in magenta, the dsRed marker in red.

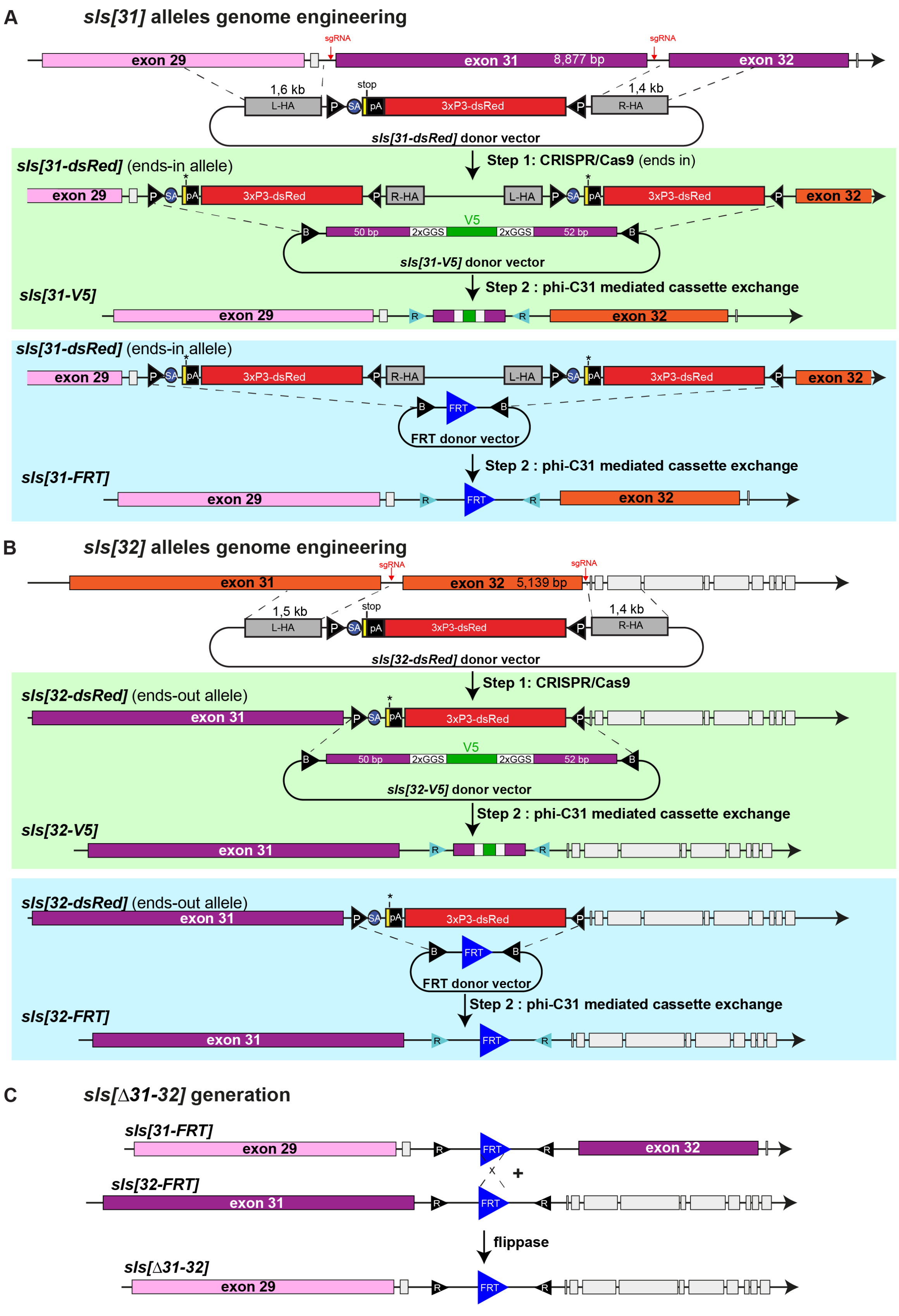
A
sls[31] alleles genome engineering
sgRNA
exon 29
exon 31
8,877 bp
exon 32
1,6 kb
stop
1,4 kb
L-HA
3xP3-dsRed
R-HA
sls[31-dsRed] donor vector
sls[31-dsRed] (ends-in allele)
Step 1: CRISPR/Cas9 (ends in)
V5
50 bp
2xGGS
52 bp
sls[31-V5] donor vector
sls[31-V5]
Step 2 : phi-C31 mediated cassette exchange
FRT donor vector
sls[31-FRT]
B
sls[32] alleles genome engineering
5,139 bp
1,5 kb
sls[32-dsRed] donor vector
sls[32-dsRed] (ends-out allele)
Step 1: CRISPR/Cas9
sls[32-V5] donor vector
sls[32-V5]
sls[32-FRT]
C
sls[Δ31-32] generation
flippase
sls[Δ31-32]

**Figure S2**

**Fig. S2 – *sls[31]* and *sls[32]* CRISPR - RMCE editing**

**(A)** Scheme showing the replacement or deletion of *sls* exon 31. Step 1: the target exon was replaced by a splice acceptor (SA)-3xstop-SV40 terminator (pA)-3xP3>dsRed cassette flanked by attP sites (P) using the CRISPR/Cas9 system ('ends-in' integration). Step 2: phi-C31-mediated cassette exchange (RMCE) was performed to replace the dsRed cassette with a V5 tag maintaining the splicing regulation of the exon 31 or an FRT site. **(B)** The same strategy was used to replace or delete *sls* exon 32 (ends-out allele). **(C)** Generation of *sls[Δ31-32]* by flippase expression (heat-shock control) in the germline of *sls[31-FRT] / sls[32-FRT]*. The successful deletion of both exons was identified by PCR.

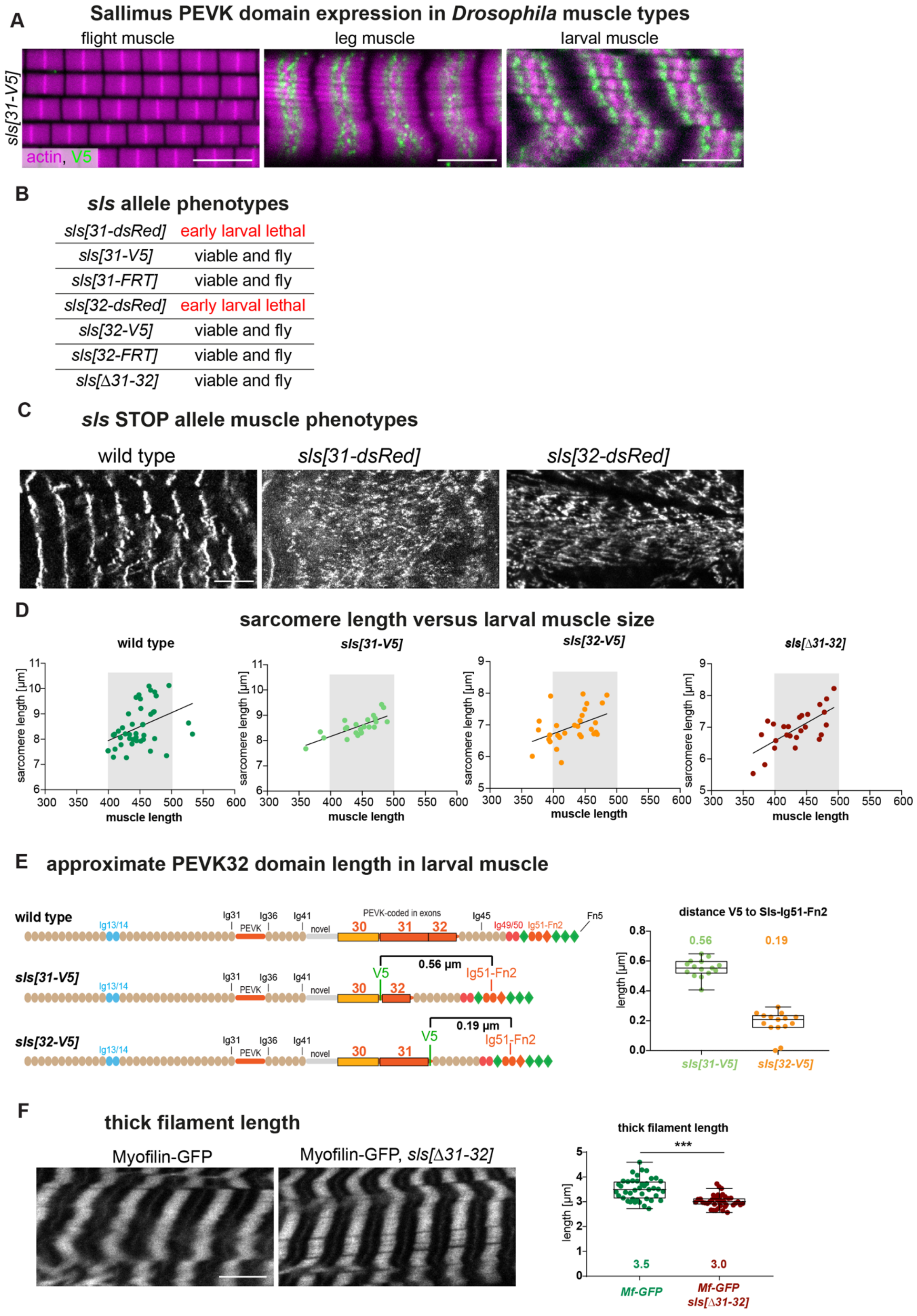
A
Sallimus PEVK domain expression in Drosophila muscle types
flight muscle
leg muscle
larval muscle
sls[31-V5]
actin, V5
B
sls allele phenotypes
sls[31-dsRed] early larval lethal
sls[31-V5] viable and fly
sls[31-FRT] viable and fly
sls[32-dsRed] early larval lethal
sls[32-V5] viable and fly
sls[32-FRT] viable and fly
sls[Δ31-32] viable and fly
C
sls STOP allele muscle phenotypes
wild type
sls[31-dsRed]
sls[32-dsRed]
D
sarcomere length versus larval muscle size
wild type
sls[31-V5]
sls[32-V5]
sls[Δ31-32]
sarcomere length [µm]
muscle length
E
approximate PEVK32 domain length in larval muscle
wild type
sls[31-V5]
sls[32-V5]
Ig13/14
Ig31
Ig36
Ig41
PEVK
novel
PEVK-coded in exons
30
31
32
Ig45
Ig49/50
Ig51-Fn2
Fn5
V5
0.56 µm
0.19 µm
distance V5 to Sls-Ig51-Fn2
length [µm]
0.56
0.19
F
thick filament length
Myofilin-GFP
Myofilin-GFP, sls[Δ31-32]
thick filament length
***
length [µm]
3.5
3.0
Mf-GFP
Mf-GFP sls[Δ31-32]

**Figure S3**

**Fig. S3 – Sls-V5 expression and *sls* alleles sarcomere phenotypes**

**(A)** Flight, leg and larval muscles of *sls[31-V5]* stained for actin (magenta) and V5 (green). Note the absence of V5 from flight muscles. Scale bars: 5 µm. **(B)** Viability of *sls* alleles. Note that *sls[31-dsRed]* and *sls[32-dsRed]* are lethal at early larval stages; the other *sls* deletion alleles are viable and can fly normally. **(C)** Stage 17 control wild-type embryo (*Zasp66-GFP / +; Df(3L)BSC366 /* +), compared to *sls[31-dsRed]* and *sls[32-dsRed]* embryos (*Zasp66-GFP / +; Df(3L)BSC366 / sls[31-dsRed]* and *Zasp66-GFP / +; Df(3L)BSC366 / sls[32-dsRed]*). Note the severely affected sarcomere pattern in the mutants. **(D)** Relation between VL3 muscle length and sarcomere length in wild type (*w[1118]*); *sls[31-V5]*; *sls[32-V5]* and *sls[Δ31-32]* third instar larvae. Muscles from 400 to 500 µm length (grey boxes) were used for quantifications in Fig. 2D. **(E)** Approximate length of Sls PEVK protein region encoded in *sls* exon 32, estimated by measuring the distance between V5 tag and Sls-Nano42 nanobody (recognizing Sls-Ig51-Fn2) in *sls[31-V5]* and *sls[32-V5]*. The difference between both indicates the PEVK32 length in larval sarcomeres. **(F)** Thick filament length by quantifying Mf-GFP signal length in wild-type *(Mf-GFP*) and *sls[Δ31-32]* VL3 larval muscles. Scale bar: 10 µm. Mann Whitney test, ***: $p< 0.001$.

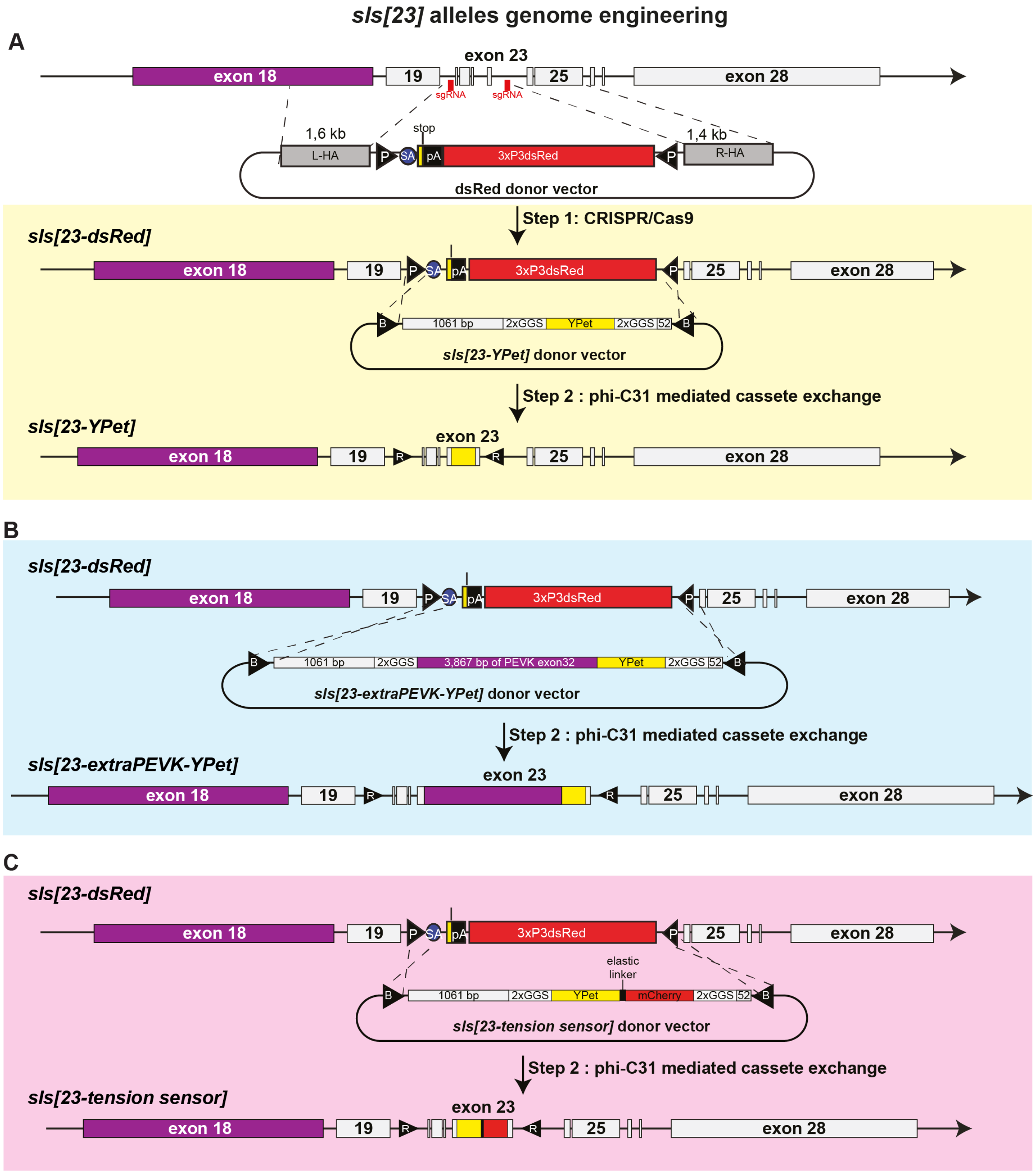


**Figure - S4**

**Fig. S4 – *sls[23]* CRISPR - RMCE editing**

**(A-C)** Schemes showing the targeting of *sls* exon 23. **Step 1:** the *sls* exons 20 to 23 were replaced by a splice acceptor (SA)-3xstop-SV40 terminator (pA)-3xP3>dsRed cassette flanked by attP sites (P) using CRISPR/Cas9. **Step 2:** phi-C31-mediated cassette exchange was performed to replace the dsRed cassette with wild-type *sls* exons 20 to 22 and *sls* exon 23 with an inserted YPet (A), or 3867 bp of *sls* exon 32 plus YPet (B) or different tension sensors (C).

A

Sound recording set up

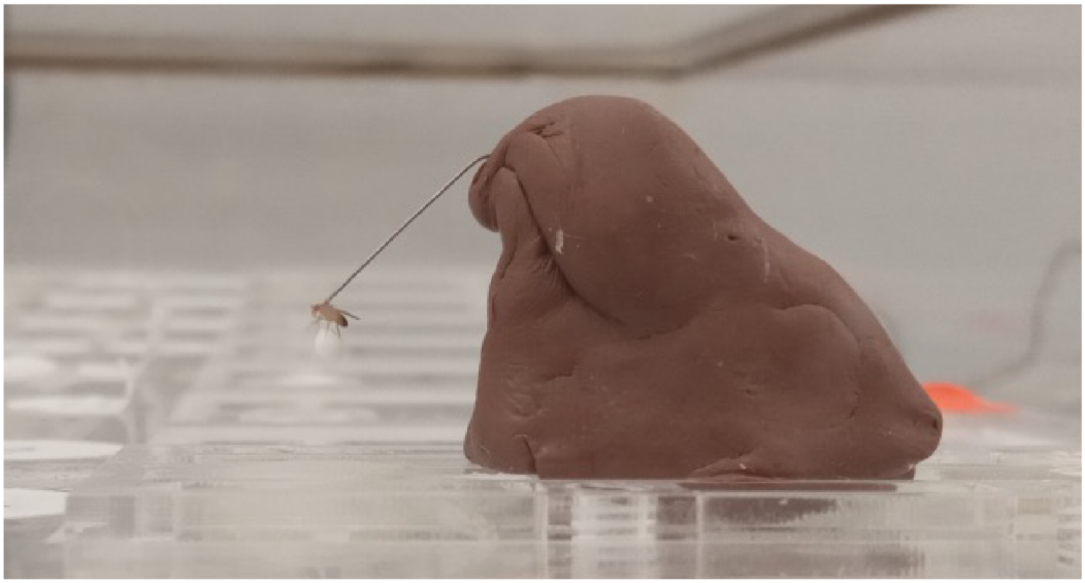

B

Individual sound traces

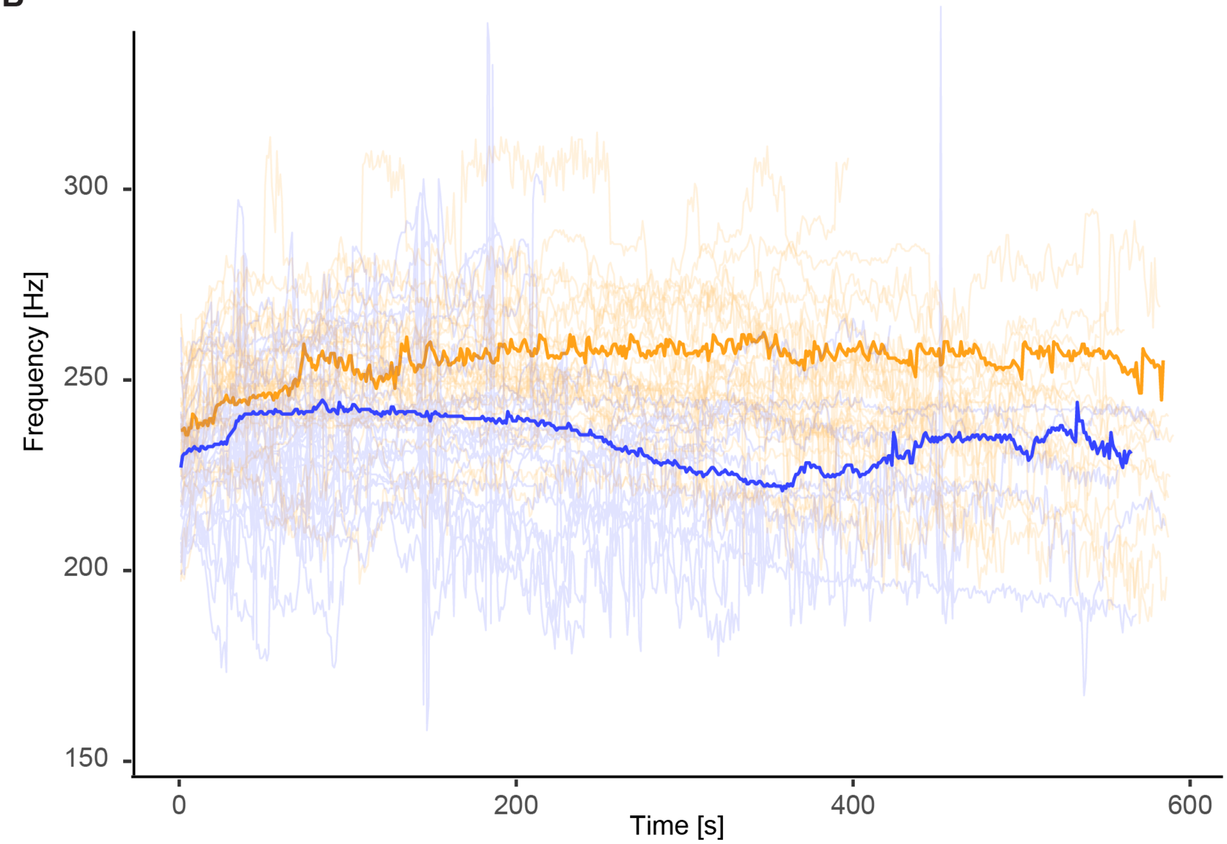


**Figure S5**

**Fig. S5 – wing beat frequency set-up and recordings**

**(A)** Wing beat sound recording set-up. The fly is tethered and rests on a paper ball. Upon removal of the ball, it starts to fly and the microphone below will record the sound frequency. (B) Individual sound traces recorded over 10 minutes. *sls[23-YPet]* plotted in orange and *sls[23-extraPEVK-YPet]* in blue. One representative trace is highlighted. Note the lower frequency and the larger variation in *sls[23-extraPEVK-YPet]*.

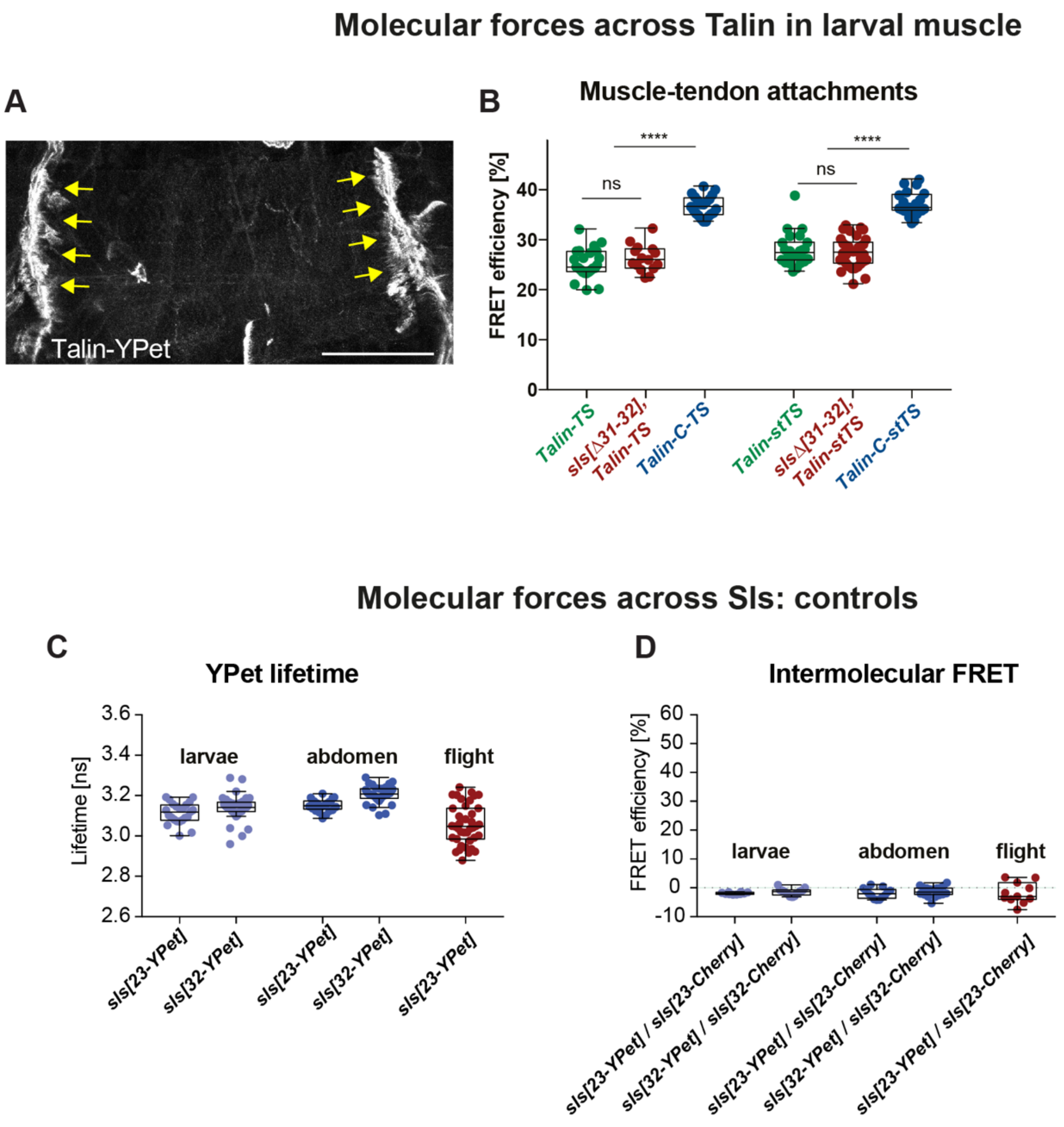


**Figure S6**

**Fig. S6 – Talin and Sallimus molecular forces**
**(A)** Expression of Talin (*rhea*)-YPet in larval muscle. Talin localises at the muscle attachments. **(B)** FLIM-based FRET quantification of Talin molecular forces comparing Talin-TS and Talin-stTS as well as C-terminal no force controls (Talin-C-TS, Talin-C-stTS) in wild type and *sls[Δ31-32]* living third instar larvae at muscle attachments. Note the reduction of FRET in Talin-TS and Talin-stTS in wild type and *sls[Δ31-32]* compared to Talin-C-TS or Talin-C-stTS. Tukey's multiple comparisons test, ns: $p>0.05$, ***: $p<0.001$. **(C)** Fluorescence lifetime (FLIM) of *sls[23-YPet]* and *sls[32-YPet]* in larval, abdominal and flight muscles. **(D)** Intermolecular FRET in larval, abdominal and flight muscles in the indicated trans-heterozygous genotypes. No intermolecular FRET is detected.

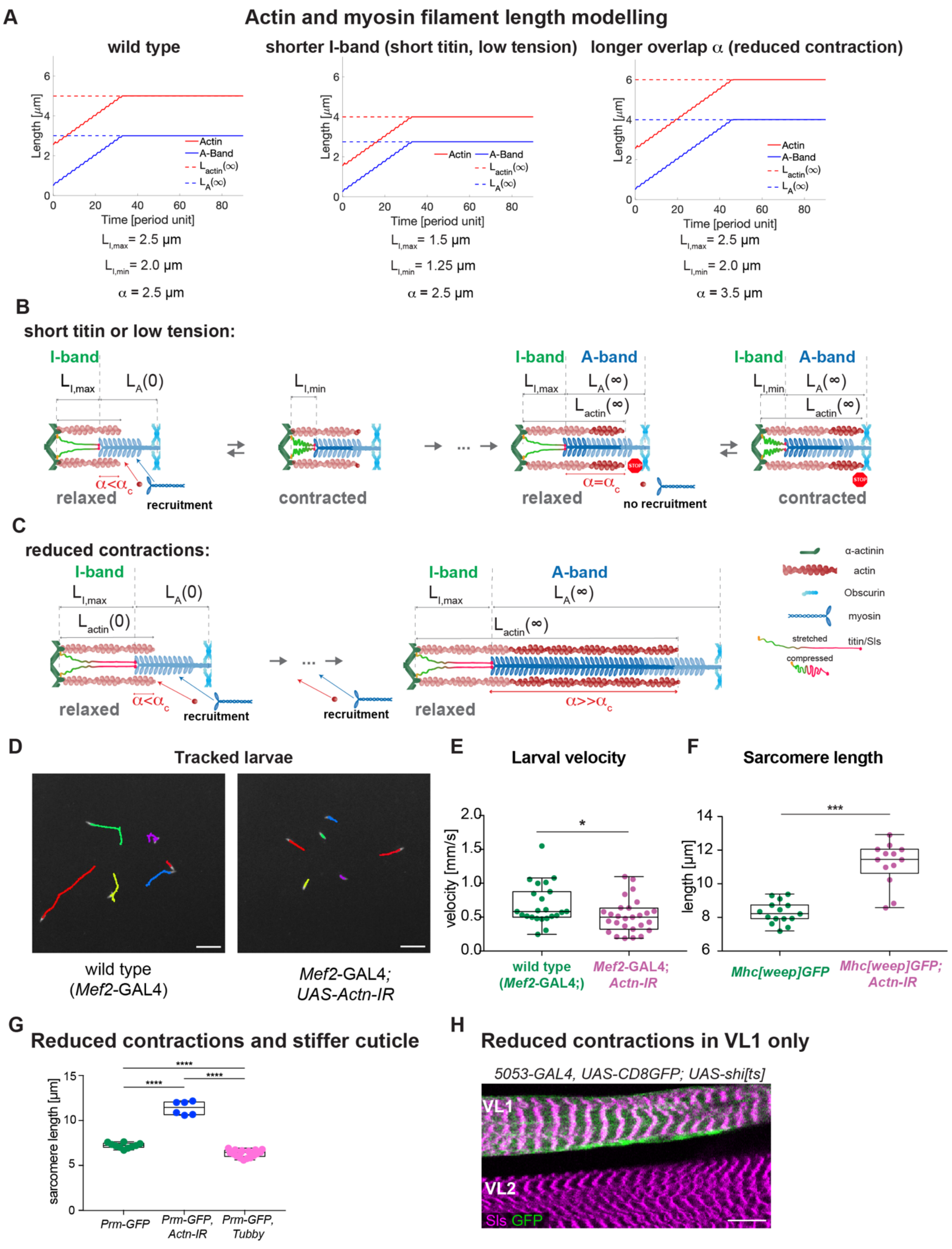
A
Actin and myosin filament length modelling
wild type
shorter I-band (short titin, low tension)
longer overlap α (reduced contraction)
Length [μm]
Time [period unit]
Actin
A-Band
$L_{actin}(\infty)$
$L_A(\infty)$
$L_{I,max}$ = 2.5 μm
$L_{I,min}$ = 2.0 μm
α = 2.5 μm
$L_{I,max}$ = 1.5 μm
$L_{I,min}$ = 1.25 μm
α = 2.5 μm
$L_{I,max}$ = 2.5 μm
$L_{I,min}$ = 2.0 μm
α = 3.5 μm
B
short titin or low tension:
I-band
A-band
$L_{I,max}$
$L_A(0)$
$L_{I,min}$
$L_A(\infty)$
$L_{actin}(\infty)$
$\alpha<\alpha_c$
$\alpha=\alpha_c$
relaxed
contracted
recruitment
no recruitment
C
reduced contractions:
$L_{actin}(0)$
$\alpha>>\alpha_c$
α-actinin
actin
Obscurin
myosin
stretched
compressed
titin/Sls
D
Tracked larvae
wild type
(*Mef2*-GAL4)
*Mef2*-GAL4;
*UAS-Actn-IR*
E
Larval velocity
velocity [mm/s]
wild type
(*Mef2*-GAL4;)
*Mef2*-GAL4;
*Actn-IR*
F
Sarcomere length
length [μm]
*Mhc[weep]GFP*
*Mhc[weep]GFP;*
*Actn-IR*
G
Reduced contractions and stiffer cuticle
sarcomere length [μm]
*Prm-GFP*
*Prm-GFP,*
*Actn-IR*
*Prm-GFP,*
*Tubby*
H
Reduced contractions in VL1 only
*5053-GAL4, UAS-CD8GFP; UAS-shi[ts]*
VL1
VL2
Sls GFP

**Figure S7**

**Fig. S7 – Actin-myosin filament length scaling**

**(A)** Modelling of the actin and myosin filament length growth over time, as predicted by the mathematical model. Note a plateau that is approached for both. Left: wild type condition with long I-band resulting in a long A-band. Middle: short I-band mutant (or low tension) results in a short A-band. Right: increased actin-myosin overlap $\alpha$ (mimicking reduced contraction) results in longer I- and A-bands and longer actomyosin overlap. **(B)** Mathematical model scheme of a short PEVK *sls* mutant or low I-band tension, which corresponds to shorter I-band lengths (both in contracted and relaxed states). When $\alpha = \alpha_C$ is reached, no new subunits are recruited and only a shorter A-band length (compared to wild type in Figure 5A) is reached. **(C)** Mathematical model scheme of a sarcomere with reduced contraction ($\alpha >> \alpha_C$) during development. Recruitment of actin and myosin subunits does not stop, hence a much longer myosin filament with a much longer actomyosin overlap (compared to wild type in Figure 5A) is generated. **(D)** Tracks of crawling larvae from Video S1. Scale bar represents 1 cm. (**E**) Quantification of larval velocity of wild-type and *Mef2*-GAL4, *Actn-IR* larvae (Mann Whitney test, *: $p < 0.05$). **(F)** Sarcomere length quantification of wild type (*Mhc[weep]GFP, Mef2*-GAL4) and *Actinin* knockdown larvae (*Mhc[weep]GFP, Mef2*-GAL4, *Actn-IR*). **(G)** Sarcomere length quantifications of the data shown in Figure 5E. **(H)** Expression of *shibire*$^{TS}$ only in VL1 muscle (marked by UAS-CD8-GFP in green) causes longer sarcomeres compared to neighbouring VL2 muscle (Z-disc labelled by Sls N-term in magenta). Scale bar is 20 µm.

**Table S1 (separate file)** – Titin evolutionary tree, species and protein names.

**Video S1 (separate file) –** Tracked wild type and *Mef2*-GAL4, *Actn-IR* larvae.

**Data S1 (separate file)** – Archive of all titin protein FASTA sequences used for evolutionary tree in Figure 1.

**Data S2 (separate file**) – Data of Figure 2

**Data S3 (separate file**) – Data of Figure 3

**Data S4 (separate file)** – Data of Figure 4

**Data S5 (separate file**) – Data of Figure 5

**File S1 (separate file**) – Python script to calculate the PEVK content in the titin sequences.
