## Supplementary material for "Titin-dependent biomechanical feedback tailors sarcomeres to specialised muscle functions in insects": File S1 - code to calculate the PEVK content: Loreau_File S1 Code_PVEK_count.py

```
#!/usr/bin/python3
# -*- coding: utf-8 -*-

## Import modules ##
import click
from Bio import SeqIO

@click.command()
@click.argument("fasta_file")
@click.argument("region_size")
def PVEK_count(fasta_file, region_size) :
	## Reading the fasta ##
	record = SeqIO.read(fasta_file, "fasta")
	## Extract the sequence ##
	sequence = record.seq
	## Name the output with the identifier of the sequence ##
	outputfile = str(record.name) + ".tsv"
	
	with open(outputfile, "w") as out:
		## Print the headers of the columns in the output ##
		print("start\tend\tproportion\tsequence", file = out)
		## The sequence is read through a window of "region_size" amino acids ##
		for i in range(0,len(sequence),int(region_size)) :
			## Counter of the P, E, V, and K ## 
			letter_count = 0
			## Extract the region of size "region_size ##
			region = sequence[i:i+int(region_size)]
			
			## For each amino acid in the region, P, E, V and K are counted ##
			for aa in region:
				if aa == "P" or aa == "E" or aa == "V" or aa == "K":
					letter_count += 1
				## Proportion of P, E, V and K per region ##
				if len(region) == region_size :
					prop = float(letter_count) / float(region_size) * 100
				else :
					prop = float(letter_count) / float(len(region)) * 100
			## Printing the start, end and proportion of the region in the output ##
			print(str(i) + "\t" + str(i+int(region_size)) + "\t" + str(prop) + "\t" + str(region), file = out)


if __name__ == "__main__":
	PVEK_count()
```
